# Large-scale functional coupling and computational modelling reveal frontotemporal hotspots in distributed network connectivity during working memory recognition, encoding, and retrieval

**DOI:** 10.64898/2026.07.23.740204

**Authors:** Wuyang Zhang, Haoru Zhang, Yuantao Deng, Xinran Deng, Yuchao Ji, Genbo Wang, Chen Yao, Yuanqing Wang, Yingchao Yu, Yiting Liu, Yuanwu Zhu, Wenbo Wang, Lanhui Huang, Sixian Li, Yue Yuan, Junfan Chen, Aoqing Luo, Yaochu Jin, Junxi Chen, Yuchen Xiao

## Abstract

Working memory depends on coordinated activity across distributed brain networks, but how these interactions support recognition, encoding, and retrieval remains unclear. Here, we analyzed intracranial local field potentials from 1,652 sites in 36 epilepsy patients performing a naturalistic card-matching task. Phase-amplitude and phase-phase coupling revealed widespread condition-dependent interactions, with frontal and temporal regions emerging as recurrent network hubs. Event-related analyses showed reproducible within-trial coupling dynamics across participants. Directed phase transfer entropy further identified frequency-specific information flow, including frontal regions acting as a theta-band source and an alpha-band sink. Finally, a time-delay-embedded hidden Markov model identified task-locked latent states with condition-dependent occupancy trajectories and distinct coherence profiles. Together, these results suggest that working memory is implemented through dynamic, directionally organized large-scale networks, in which frontotemporal hubs coordinate flexible transitions among distributed neural states rather than operating in isolation.

## Introduction

Working memory (WM) constitutes core mental faculties of the human brain, characterized by the retention and manipulation of a limited amount of information over a short period of time. Real-world WM processes can be highly dynamic, flexible, and complex^1,2^, thus requiring intricate orchestration from multiple brain regions and precise spatiotemporal coordination among neural assemblies. Working memory impairments have been associated with various pathological conditions including Alzheimer’s disease^3^ and schizophrenia^4^, for which there are few efficacious treatments to date, calling for further mechanistic interrogation at macroscopic, mesoscopic, and microscopic levels.

Despite a large number of studies, the anatomical basis for working memory is still in debate. Accumulating studies with heterogeneous results only seem to obscure rather than resolve the situation. Multiple studies have propounded important roles of the medial temporal lobe (MTL) in WM^5-7^; however, the famous case of patient H.M. presents as an opposing evidence, where bilateral resection of the MTL only abolished the ability to form long-term memory while working memory was retained^8,9^, although one may argue that tissues “escaped” from lobectomy could do the work. Furthermore, lesions in non-MTL regions, such as the temporal-parietal junction, specifically obliterated short-term rather than long-term memory^8^. In accordance with the “central executive” component in Baddeley’s model of working memory^8^, the prefrontal cortex has been shown to be activated in working memory tasks^10,11^. As the candidates for the neuroanatomical location of WM keep expanding and the localizationist view of WM becomes less tenable, the notion of “distributed network” offers plausible alternative accounts, from extreme propositions assuming omnipotency of the neocortex^12^ to moderate ones that bridge with localization-of-function by highlighting a few significant regions among large-scale networks^13,14^. Insofar as the seemingly opposing views haven’t fought out a point, direct neurobiological evidence from reliable measurements and unbiased analyses would be propitious to advancing the field.

In this work, we recorded human intracranial local field potentials (LFP) from 36 pharmacologically-intractable epilepsy patients who received electrode implantation for seizure monitoring. Local field potentials measured from thin (0.8 mm in diameter) stereo-EEG electrodes yield high spatial and temporal resolution, and thus high signal-to-noise ratio and data precision. Subjects completed a card-matching working memory game which is designed to generate high levels of flexibility, load, dynamics, and autonomy to mimic real-world situations. Unlike most human intracranial studies, we forfeited the approach to focusing on one or two selected anatomical regions yet unbiasedly investigated our entire sample of electrode locations. Through this methodology, the relative contributions from different regions to working memory processes could be reliably weighed, the WM “hotspots” be confirmed or rejected, and the intricate connections in distributed networks be revealed.

In previous work^2^, we focused on single-electrode analysis to investigate the role of gamma oscillations in working memory processes. For the current study, we explored both within- and cross-regional functional coupling (FC), both in the same frequency band and between differential frequencies. In electrophysiological studies, functional coupling refers to patterned interactions between spectral features of neural oscillations, whose putative functions include neural representation, local computation, long-range communication, and internal clocking^15-17^. Given these fundamental neural processes, FC is thought to contribute to numerous cognitive functions^18^. Pertaining to working memory, research indicates that FC is involved in its representation^19,20^, maintenance^21,22^, and executive control^19^.

The significance of studying functional coupling lies in its capacity to reflect the mechanisms of information transfer and exchange across different spatial and temporal scales within neurophysiological activities, as measured from LFP, scalp-EEG, and MEG. Recent perspectives on working memory^8^ imply the requirement of effective interaction across regions and suggest distinct WM processes^23,24^. Accordingly, we separated WM processes into recognition, encoding, and retrieval, and investigated the functional coupling in forms of phase-amplitude coupling (PAC) and phase-phase coupling (PPC) within each memory stage. Following the same logic as we investigated all anatomical regions, we considered all frequencies for within-frequency analysis and all biologically plausible frequency combinations for cross-frequency analysis. Most previous studies on neural oscillations pre-selected only one or two frequencies a priori, most likely due to computational unwieldiness and interpretative challenge. Although results from this approach may appear clean and lucid in isolation, it is difficult to integrate these discrete pieces of information into a coherent and informative neurobiological model.

To extract insightful results from the massive amount of data out of regionally and spectrally inclusive approaches, we developed a systematic and generalizable pipeline for large-scale connectivity analysis using LFP signals. Regional importance was determined in a “gradually zooming in” manner, from macroscopic (lobes or large structures), to mesoscopic (finer regions), and finally microscopic (electrodes). Weighted connectivity was then constructed both to verify the regional importance and to infer connecting direction and strength. K-means clustering for time series combined with dynamic time warping was leveraged to pinpoint FC patterns that were common across individuals.

Together, our results reveal a distributed but frontotemporal-centered architecture for human working memory. Across PAC, PPC and CCI analyses, frontal regions—particularly prefrontal/orbitofrontal control areas—and temporal regions, including lateral temporal and medial temporal structures, emerged as recurrent high-contribution nodes rather than isolated anatomical loci. Directed phase transfer entropy (dPTE) further showed that these hubs participate in frequency-specific directional communication, with frontal regions tending to act as theta-band information sources while showing more sink-like or gating roles in alpha-band interactions. Finally, a time-delay-embedded hidden Markov model (TDE-HMM) identified a low-dimensional repertoire of task-locked network trajectories, including a crossover motif between early- and late-dominant states and a near-stationary motif with stable state occupation. State-resolved coherence analyses linked these latent trajectories to complementary memory-related network modes, characterized by frequency-specific reweighting of frontotemporal and broader distributed pathways. By integrating large-scale functional coupling, directional information flow and latent-state modeling, our study reframes human working memory as a dynamic communication process in distributed networks, providing constraints for computational theories and potential targets for memory-oriented neuromodulation.

## Results

Intracranial LFPs were recorded from 1652 electrodes (**Table S1**) in 36 subjects (**Table S2**) who completed a card-matching memory task, as described in our previous study^2^ (**Figure 1, Movie S1**). Subjects were instructed to locate all pairs of images in a tile array that was initially masked. In each trial, two tiles were revealed sequentially in a self-paced manner. A match was achieved if the two tiles contained the same image; otherwise, a mismatch occurred. Already matched tiles were filled with green color and could not be clicked again. The level of difficulty increased from 3 × 3 to 7 × 7 arrays and remained at 7 × 7 until the end of the session.

**Figure 1.**
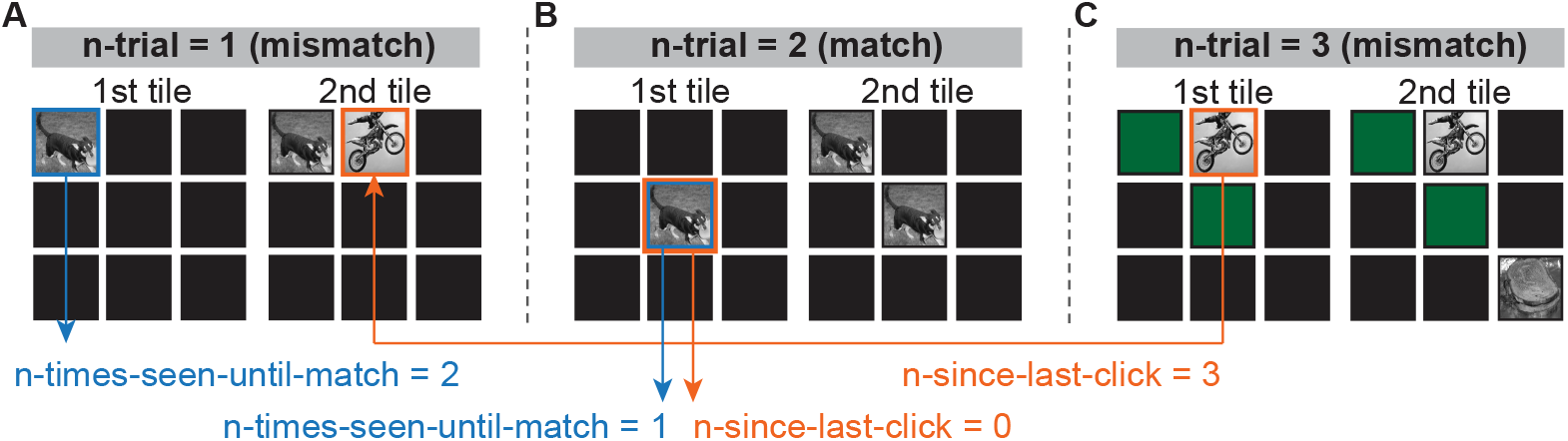
Experimental paradigm. **A-C.** Illustration of three consecutive trials in a 3 × 3 board. In each trial, two tiles were sequentially clicked and immediately revealed. Tile 1 remained “on” while waiting for the second click. Once the second tile was flipped, subjects were given 1 second to view the result—either a match or a mismatch—followed by both tiles turning green and frozen (if a match) or both tiles being re-masked (if a mismatch). N-times-seen-until-match: the total number of flips for a given tile before it was successfully matched in a block. N-since-last-click: the number of flips elapsed since the current tile was flipped previously.

### Widely distributed networks during working memory processes

Working memory (WM) is not localized to a single anatomical region or to isolated neural modules. Instead, it recruits distributed networks that involve sensory and association cortices as well as subcortical structures, where precise temporal and spatial coordination is essential for task execution. We therefore first investigated phase-amplitude coupling (PAC), the coupling between low-frequency phase and high-frequency amplitude, in all single electrodes and all simultaneously recorded pairs of electrodes, yielding both within-regional and cross-regional functional coupling. PAC strength was calculated as the Modulation Index (MI, Methods), and statistical inference incorporated surrogate-based significance assessment and false-discovery-rate (FDR) correction where appropriate.

All FC analyses were restricted to neural signals between tile 1 and tile 2. We did not consider activity after tile 2 because the second flip was followed by strong feedback and decision-related effects once subjects realized whether the trial was a success (match) or a failure (mismatch). Furthermore, activity after the second flip was a complex mixture of responses to two tiles with the same or different images at different screen locations, making it difficult to disentangle and interpret. For LFP signals sandwiched between two flips in each trial, we defined conditions under distinct WM processes based on behavioral metrics (Methods). For recognition, we contrasted novel (first-seen) versus old tiles. For encoding, we considered first-seen tiles only and separated them into better encoded (BE) and worse encoded (WE) items using the parameter n-times-seen-until-match (ntsum, **Figure 1**); BE corresponds to the smaller ntsum cluster, whereas WE corresponds to the larger ntsum cluster. To redress potential subject variability in memory encoding capacity, unsupervised k-means clustering was performed for each subject to yield a personalized ntsum threshold. For retrieval, we compared match versus mismatch trials after removing random-match trials and excuse-mismatch trials (**Methods**).

We first compared PAC under each set of conditions during recognition, encoding and retrieval, respectively. According to the putative biological and computational mechanisms of phase-amplitude coupling, not all combinations of the frequency for phase and the frequency for amplitude yield meaningful insights into neural functional coupling^25,26^. For example, low gamma-high gamma PAC is highly correlated with muscle artifacts, and alpha-beta PAC may contain artifactual harmonic coupling^25^. In addition, oscillations slower than theta were not analyzed because the relatively short tile1-to-tile2 interval limited reliable estimation of very slow rhythms^27^. Based on these considerations, we focused on four PAC frequency combinations: theta-low gamma, theta-high gamma, alpha-low gamma and alpha-high gamma. For each electrode pair, condition and PAC frequency combination, a surrogate distribution was generated, and only electrode-pair entries showing above-chance PAC in at least one condition were retained for population-level analysis (empirical p < 0.05). The MI values from the retained electrode pairs were then grouped by directed region pairs, where the first region indicates the phase-providing region and the second region indicates the amplitude-providing region. Condition differences in MI were assessed at the region-pair level using paired Wilcoxon signed-rank tests, followed by FDR correction across region-pair comparisons. **Figure 2C-E** shows alpha-low gamma PAC across all retained electrode pairs from all subjects. These analyses revealed non-confined, widespread differences in PAC strength between opposite conditions in each WM stage, indicating broadly distributed functional coupling during recognition, encoding and retrieval. Other frequency combinations showed similar distributed effects (**Figure S1**).

**Figure 2.**
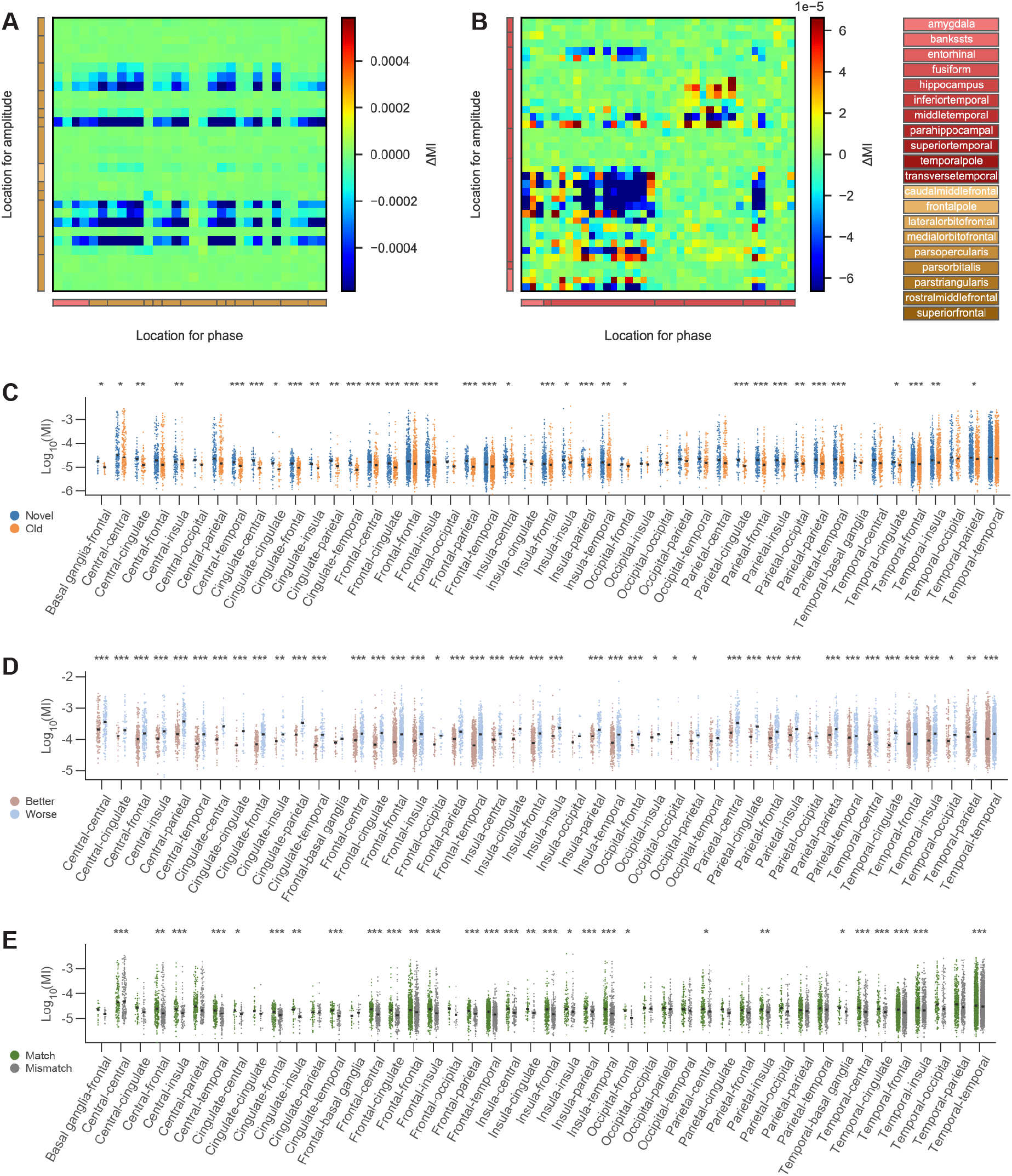
Widely distributed phase-amplitude coupling under different WM processes. **A-B.** The two panels illustrate the difference in Phase-Amplitude Coupling (**A**) between the Match and Mismatch conditions and (**B**) between the Novel and Familiar conditions for two distinct examples (e.g., two different subjects). Specifically, (**A**) displays the coupling differences for theta phase and high gamma amplitude, whereas (**B**) illustrates the differences for alpha phase and low gamma amplitude across various brain regions. The horizontal axis represents the anatomical locations providing the phase component (Location for phase), while the vertical axis lists the locations providing the amplitude component (Location for amplitude). These electrodes are categorized by their general anatomical regions (Frontal, Temporal, Parietal, Central, Insula, Occipital, and Cingulate) and sorted alphabetically. The color intensity in the heatmaps denotes the change in Modulation Index (ΔMI), with the value range indicated by the colorbar on the right of each panel, reflecting the difference in coupling strength between the two behavioral conditions for each electrode pair. **C**-**E**. Alpha-low gamma PAC strength in each combination of general regions from all subjects during recognition (**C**), encoding (**D**), and retrieval (**E**). Modulation index (MI) is shown in log10 scale to account for its exponential distribution. Black horizontal bars indicate the median PAC value in each condition. Location combinations containing fewer than 20 samples were removed from analysis. The asterisks highlight region pairs with statistically significant differences between conditions (Wilcoxon signed-rank test, alpha = 0.05, *p < 0.05, **p < 0.01, and ***p < 0.001). All p-values were corrected for multiple comparisons using the false discovery rate (FDR) method.

Next, we investigated the profile of phase-phase coupling during each memory process. The strength of phase-phase coupling was estimated using the phase-locking value (PLV, Methods), which reflects the consistency of phase information between two signals. By definition, only cross-electrode PPC could be computed. Analogously, for each electrode pair, condition and PPC frequency band, a surrogate PLV distribution was generated to retain only above-chance inter-electrode phase locking before population-level analysis (empirical p < 0.05 in at least one condition). Retained electrode pairs were then grouped into undirected region pairs, because PPC does not assign a phase-providing and amplitude-providing direction as PAC does. Condition differences in PLV were assessed at the region-pair level using paired Wilcoxon signed-rank tests followed by FDR correction across region-pair comparisons. **Figure 3** displays theta phase-phase coupling strength under different conditions during each WM component. A large number of region pairs demonstrated significant PPC differences between viewing novel and old stimuli (**Figure 3C**). Similarly, during WM encoding, significant PPC differences between BE and WE items were widely distributed (**Figure 3D**). In contrast, phase coherence appeared more selectively organized during WM retrieval, where frontal-frontal and frontal-temporal PPC exhibited particularly prominent differences between match and mismatch trials (**Figure 3E**). Results of other frequency bands are shown in **Figure S1-3**.

**Figure 3.**
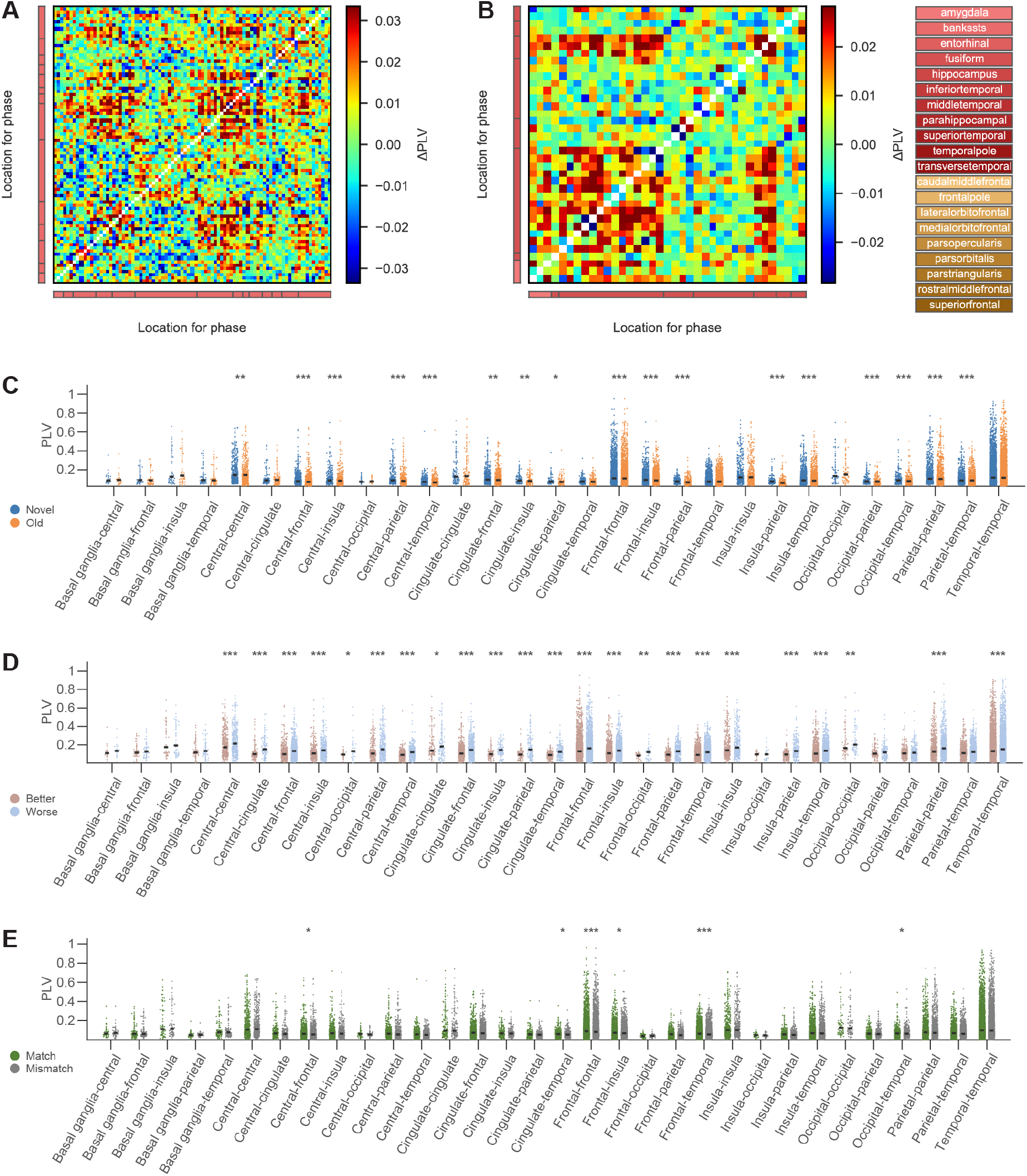
Phase-phase coupling under different WM processes. A-B. The two panels illustrate the difference in Phase-phase Coupling (**A**) of the alpha band between the Novel and Familiar conditions and (**B**) of the beta band between the Match and Mismatch conditions for two distinct examples (e.g., two different subjects). The horizontal and the vertical axes represent the anatomical locations providing the phase component. These electrodes are categorized by their general anatomical regions (Frontal, Temporal, Parietal, Central, Insula, Occipital, and Cingulate) and sorted alphabetically. The color intensity in the heatmaps denotes the change in Phase Locking Value (ΔPLV), with the value range indicated by the colorbar, reflecting the difference in coupling strength between the two behavioral conditions for each electrode pair. **C**-**E**. Theta phase-phase coupling (PPC) strength in each combination of general regions from all subjects during recognition (**C**), encoding (**D**), and retrieval (**E**). PPC strength was calculated as phase-locking value (PLV). Black horizontal bars indicate the median PLV value in each condition. Location combinations containing fewer than 20 samples were removed from analysis. The asterisks highlight region pairs with statistically significant differences between conditions (Wilcoxon signed-rank test, alpha = 0.05, *p < 0.05, **p < 0.01, and ***p < 0.001). All p-values were corrected for multiple comparisons using the false discovery rate (FDR) method.

### Frontotemporal hubs in functional coupling networks during working memory processes

The population-level PAC and PPC analyses showed that many brain regions demonstrated significant FC differences between contrastive conditions under each working memory component. Next, to facilitate direct comparison of the contribution of each region to the FC networks, we implemented a hierarchical normalization procedure - the connectivity contributive index (CCI) - which preserves the original proportional relationship among regions while accounting for the unbalanced distribution of electrode locations across subjects (Methods), a typical circumstance of human sEEG studies. A CCI of 1 indicates the highest relative contribution in a given analytical context; CCI was used as a continuous contribution index rather than as a hard threshold for classifying regions.

Figure 4 shows CCI results in alpha-low gamma PAC, theta-high gamma PAC, and an overall view combining all PAC frequency pairs. **Figure 5** displays CCI in theta and alpha phase networks, as well as the overall pattern considering all PPC frequencies. Results from other PAC frequency combinations and PPC frequencies are shown in **Figure S7** and **Figure S8.** These analyses indicated that frontal and temporal cortices played a central role in large-scale functional coupling networks during WM processes, with the frontal lobe featuring most prominently across several contexts (**Figures 4-5**). Other regions that were strongly involved in the FC networks, although less consistently than frontal and temporal lobes, included the parietal lobe and the insula (**Figures S7** and **S8**).

**Figure 4.**
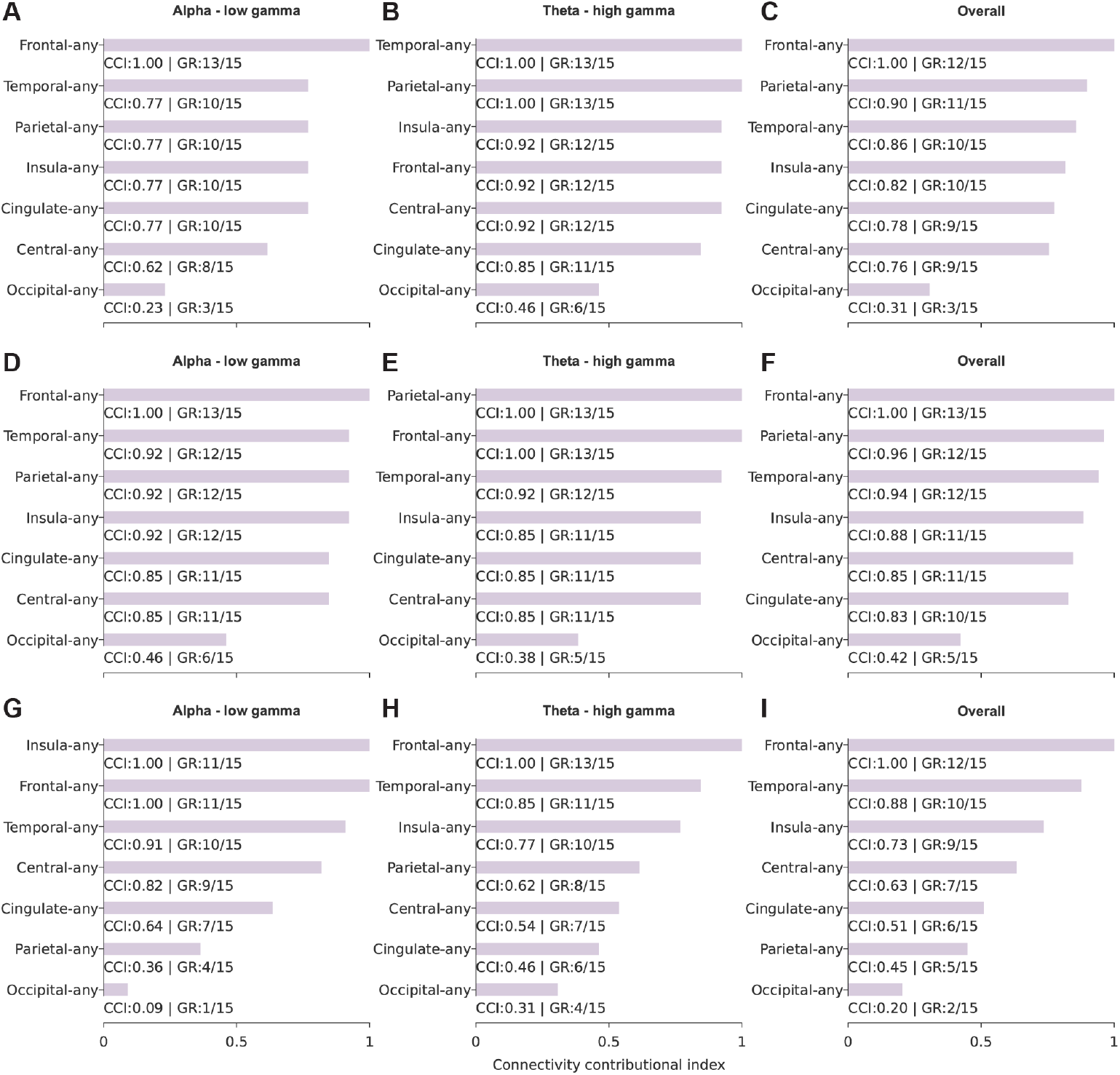
Regional contribution to PAC network. The contribution of any general region to the overall PAC network was estimated using connectivity contributive index (CCI). Each row shows CCI results of each WM process. **A**-**C**: recognition; **D**-**F**: encoding; **G**-**I**: retrieval. The left column displays results in alpha-low gamma PAC and the middle column theta-high gamma PAC. Data from other combinations are shown in **Figure S7**. The right column shows the overall CCI results combining all frequency pairs. In each panel, horizontal bar length represents the CCI value, also labelled under each bar. A CCI = 1 indicates the highest possible contribution of any region in this dataset, within each frequency combination (left/middle columns) or across all frequency combinations (right column). Region for phase and region for amplitude were not differentiated here. For example, “parietal-any” incorporated all significant PAC region pairs as long as parietal cortex was involved, irrespective of whether it provided phase or amplitude information. “GR” specifies the number of subjects who had the region as a significant PAC contributor during a certain memory process, divided by the total number of subjects who had electrodes in that general region.

**Figure 5.**
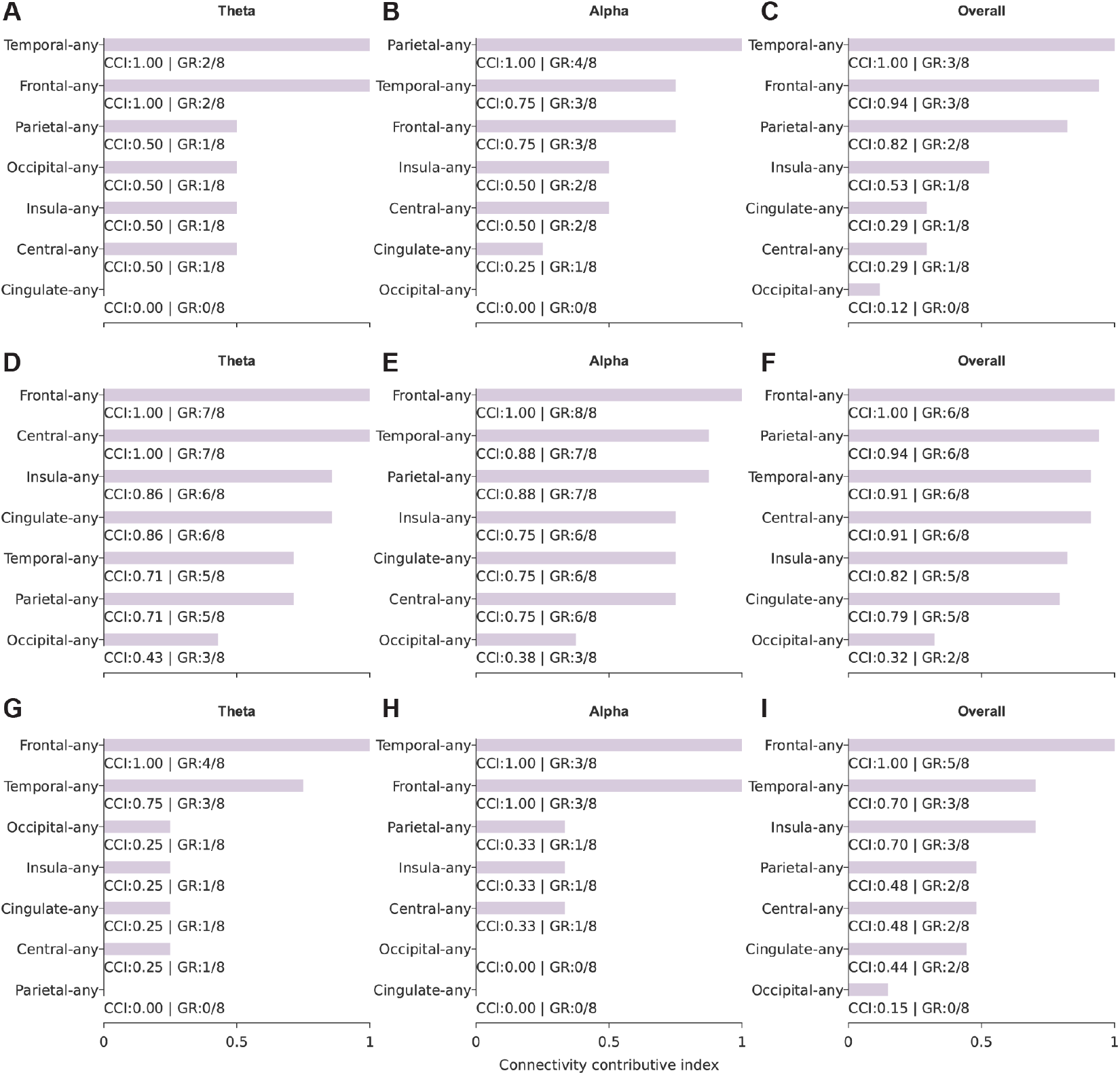
Regional contribution to PPC network. The contribution of any general region to the overall PPC network was estimated using connectivity contributive index (CCI). Each row shows CCI results of each WM process. **A**-**C**: recognition; **D**-**F**: encoding; **G**-**I**: retrieval. The left column displays results in theta PPC and the middle column alpha PP**C**. Data from other frequency bands are shown in **Figure S8**. The right column shows the overall CCI results combining all frequency bands. In each panel, horizontal bar length represents the CCI value, also labelled under each bar. A CCI = 1 indicates the highest possible contribution of any region in this dataset, within each frequency band (left/middle columns) or across all frequency bands (right column). “GR” specifies the number of subjects who had the region as a significant PPC contributor during a certain memory process, divided by the total number of subjects who had electrodes in that general region.

### Consistent event-related functional coupling dynamics

Having established that PAC and PPC differences were distributed across brain regions and centered on frontotemporal networks, we next asked whether these significant connections exhibited reproducible within-trial temporal dynamics. We examined event-related PAC (ERPAC) and event-related PPC (ERPPC) within the tile1-to-tile2 interval (**Methods**). ERPAC or ERPPC were calculated only for region pairs that demonstrated both significant condition differences in the corresponding PAC/PPC analysis and significantly higher MI or PLV than surrogate distributions in at least one condition during each WM process. This two-step selection ensured that the dynamic analyses focused on physiologically detectable and task-relevant coupling effects rather than arbitrary electrode pairs.

To examine whether there were consistent ERPAC or ERPPC trends across subjects, we employed K-means clustering for time series with dynamic time warping (DTW) as the distance metric. DTW was used to capture shared temporal structures while minimizing the impact of slight and non-critical temporal shifts in the signals. For each eligible region pair and frequency band, coupling trajectories were normalized and clustered separately within each condition, yielding a small set of representative temporal motifs.

ERPAC analyses revealed structured temporal motifs across the three WM components. As an illustrative frontotemporal/medial-temporal example, theta-high gamma ERPAC between hippocampus and middle temporal cortex was grouped into distinct clusters during recognition, encoding, and retrieval (**Figure 6**). The presence of similar motif families across novel, old, BE, WE, match, and mismatch trials suggested that PAC differences were not merely static offsets in MI, but reflected condition-dependent changes in the temporal expression of cross-frequency coupling. In parallel, ERPPC analyses identified reproducible theta-band phase-synchrony trajectories, exemplified by amygdala-lateral orbitofrontal coupling (**Figure 7**). These results indicate that the frontotemporal network identified by the static PAC/PPC and CCI analyses also exhibits temporally structured, event-related coupling dynamics.

**Figure 6.**
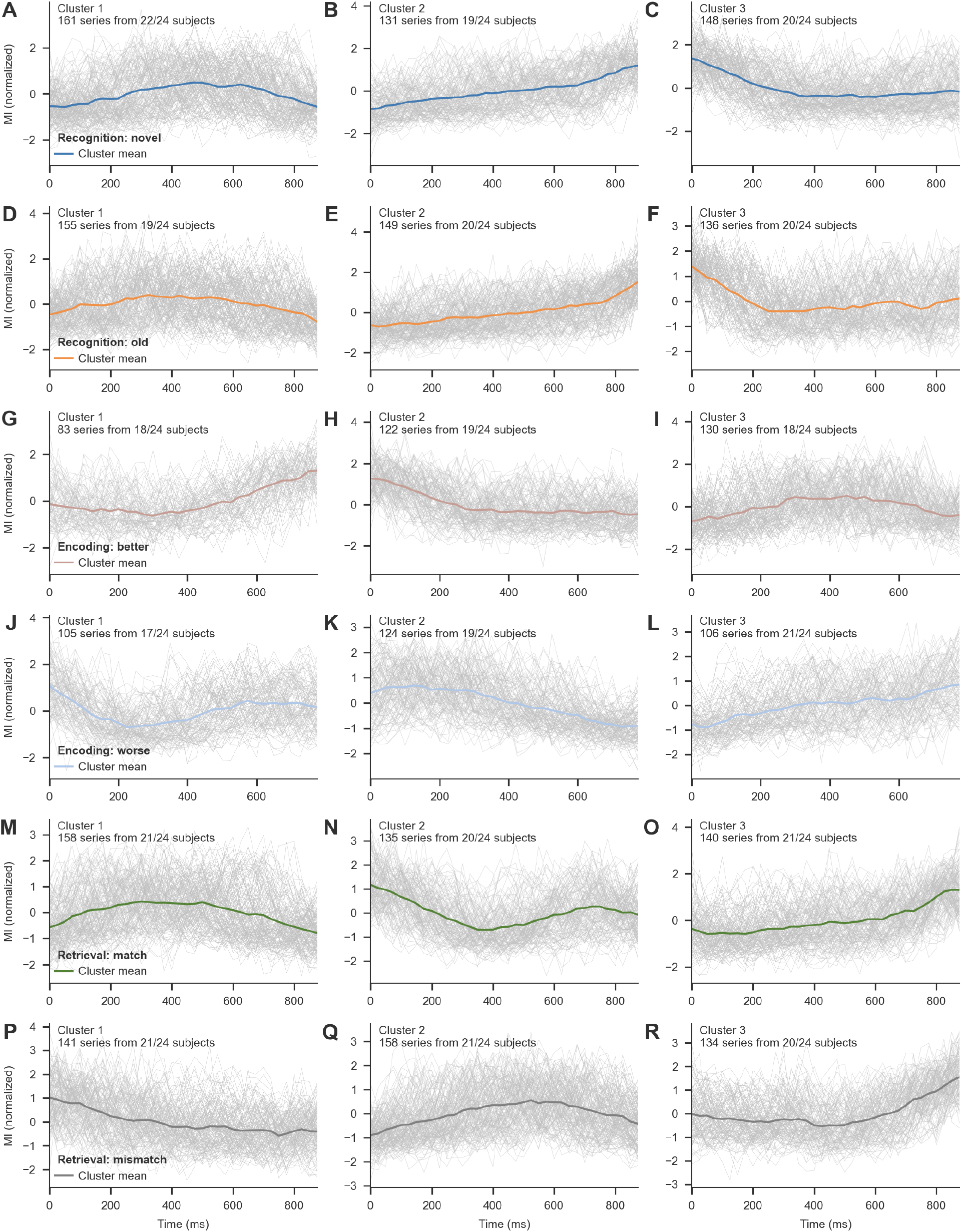
Event-related phase-amplitude coupling (ERPAC) dynamics. **A-R.** ERPAC between hippocampus and middle temporal cortex in the theta-high gamma frequency combination across three memory processes. **A-C**. Novel trials during recognition. **D-F**. Old trials during recognition. **G-I**. BE trials during encoding. **J-L**. WE trials during encoding. **M-O**. Match trials during retrieval. **P-R**. Mismatch trials during retrieval. Each panel shows one of three clusters identified by time-series K-means clustering using DTW distance. Individual time series are shown in light gray, and cluster means are shown in colored lines. The modulation index was normalized using mean-variance scaling and plotted as a function of time relative to stimulus onset. The number of time series and subjects contributing to each cluster are indicated in each panel.

**Figure 7.**
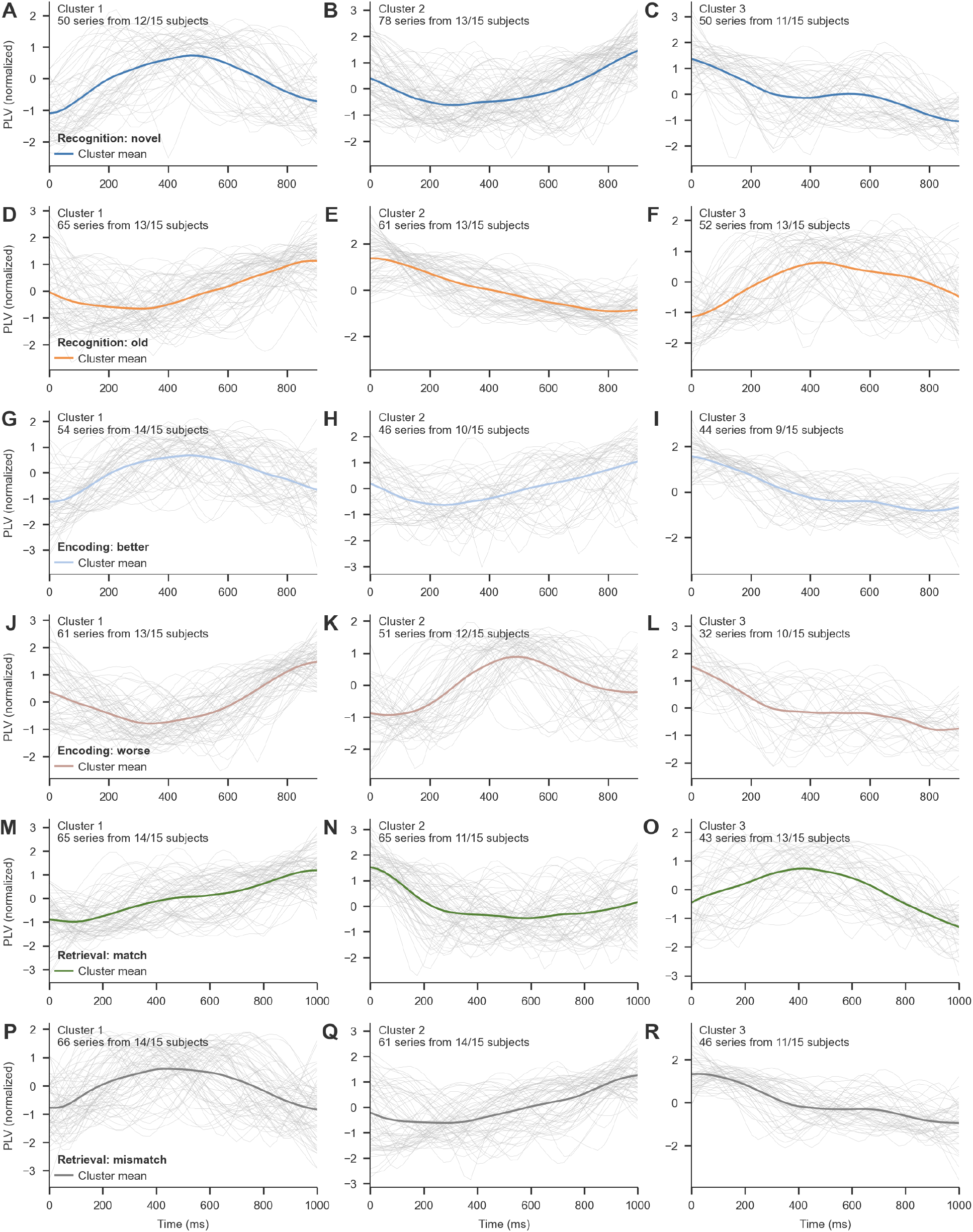
Event-related phase-phase coupling (ERPPC) dynamics. **A-R.** ERPPC between amygdala and lateral orbitofrontal cortex in the theta band across three memory processes. **A-C**. Novel trials during recognition. **D-F**. Old trials during recognition. **G-I**. BE trials during encoding. **J-L**. WE trials during encoding. **M-O**. Match trials during retrieval. **P-R**. Mismatch trials during retrieval. Each panel shows one of three clusters identified by time-series K-means clustering using DTW distance. Individual time series are shown in light gray, and cluster means are shown in colored lines. PLV was normalized using mean-variance scaling and plotted as a function of time relative to stimulus onset. The number of time series and subjects contributing to each cluster are indicated in each panel.

We further summarized the cross-subject coverage of ERPAC and ERPPC clusters across frequency bands and WM processes. Overall, no dominant frequency band or method showed a systematic coverage advantage that would trivially explain the observed temporal motifs. Instead, most clusters were represented by moderate-to-high subject coverage across recognition, encoding, and retrieval (**Figure S13**), supporting the interpretation that event-related FC dynamics were shared across participants rather than being driven by isolated individual cases.

### Directionally organized information flow across lobes

PAC and PPC quantify the strength of cross-frequency or same-frequency coupling, but they do not by themselves determine whether one region tends to lead or drive another. We therefore used directed phase transfer entropy (dPTE) to test whether the frontotemporal functional coupling networks were accompanied by directionally organized information flow (**Methods**). For each canonical frequency band, dPTE was summarized across lobe pairs and compared between the relevant conditions in recognition (novel vs old), encoding (BE vs WE), and retrieval (match vs mismatch). Directional values above or below 0.5 indicated preferential information flow in opposite directions, and condition differences were assessed at the lobe-pair level.

Multiple lobe pairs showed significant condition-dependent difference in dPTEs (Wilcoxon signed rank test, novel vs. old, better vs. worse encoded, match vs. mismatch) in each frequency band (**Figure 8B, Figure S14-S16**). All the lobe pairs had different dPTEs between match and mismatch conditions in low-gamma band (**Figure S16D**), which also had the largest proportion of channel pairs with significant different dPTEs between the two conditions (**Figure 8A**). Frontal and temporal lobes showed a strong connectivity with other lobes, which is consistent across subjects (**Figure 8A**).

**Figure 8.**
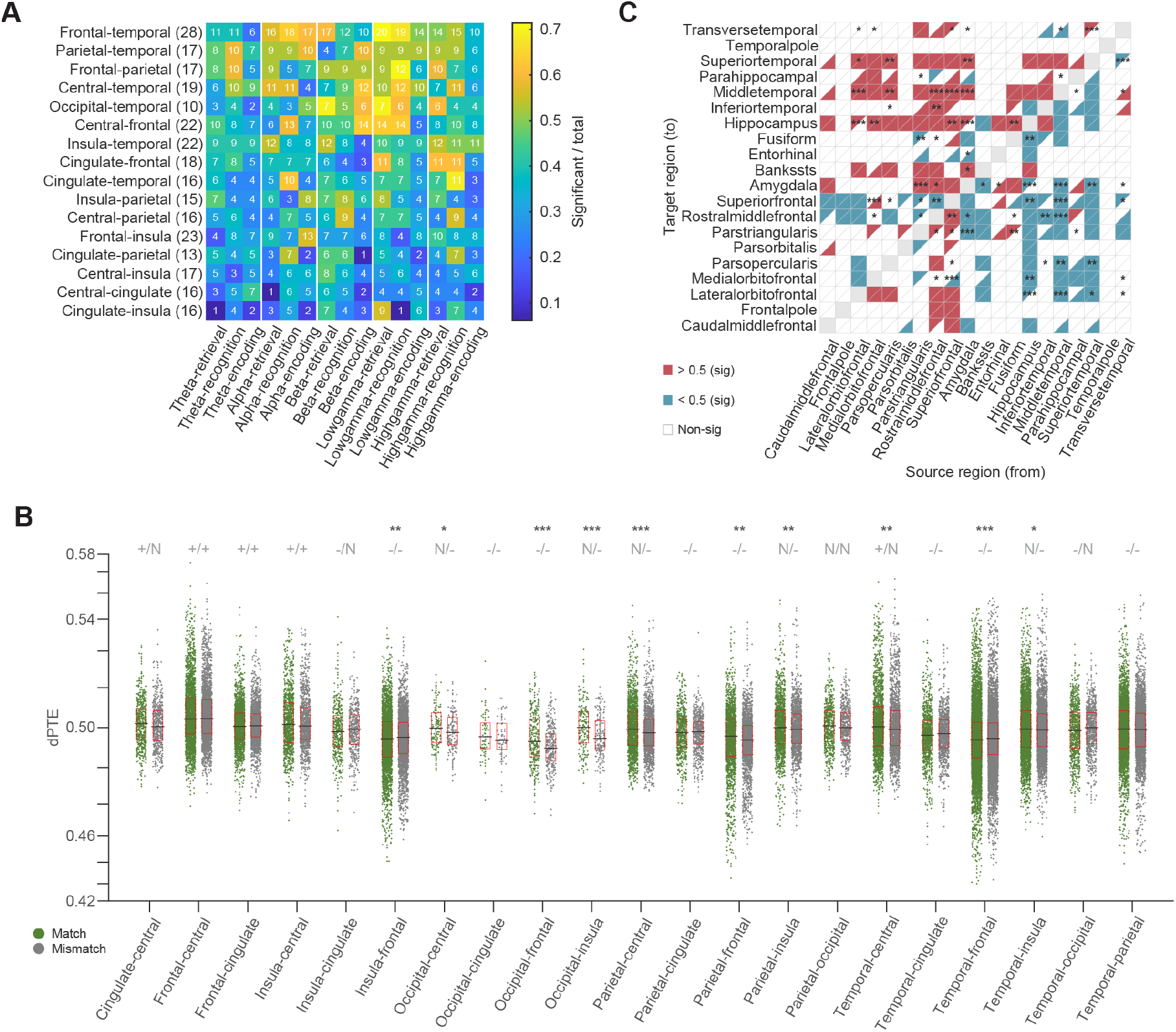
dPTEs difference among frequency bands and conditions. **A.** Numbers of subjects that showed significant condition-dependent difference in dPTEs in each frequency band. The number of subjects that have this lobe recorded shows in parenthesis. Only pairs that were recorded from over 10 subjects were displayed. **B**. dPTE in theta band across all subjects (lobe level, match vs mismatch). Each dot (green: match, gray: mismatch) corresponds to the median dPTE of a channel pair in this lobe pair. Asterisk: median dPTEs showed significant difference (Wilcoxon signed rank test, *: p < 0.05, **: p < 0.01, ***: p < 0.001) between match and mismatch conditions. + means median dPTE was significantly larger than 0.5 (Wilcoxon signed rank test, p < 0.05), while – means smaller than 0.5, and N means no significant different compared to 0.5. Black horizontal line: median dPTE of the lobe pair under each condition. Red box: interquartile range (25^th^∼75^th^ percentile) of the dPTE distribution for each condition. **C**. dPTE difference in theta band for region pairs in frontal and temporal lobes between match and mismatch conditions. Top-right and bottom-left triangles show whether the dPTEs were significantly larger (red) or lower (blue) than 0.5 under match and mismatch conditions respectively (Wilcoxon signed rank test, p < 0.05). Asterisk: median dPTEs showed significant difference (Wilcoxon signed rank test, *: p < 0.05, **: p < 0.01, ***: p < 0.001) between match and mismatch conditions.

To further investigate which region played a role in conveying different information between conditions in low-gamma band, **Figure 8C** exhibits the region pairs from frontal and temporal lobes. It was found that the dPTEs from parstriangularis (frontal lobe) to several temporal regions were significant different between match and mismatch conditions.

To confirm whether there existed information sources and sinks during the memory-related process, we used the dPTEs as weights to construct directed weighted connectivity matrix across the lobes (**Methods**). Frontal lobe showed converse patterns between theta and alpha bands. It was found that frontal lobe tended to be the information source (Net flow > 0) under theta band, while it acted as the information sink (Net flow < 0) in alpha band (16/32 subjects both in match and mismatch trials, chance level = 0.252) (**Figure 9**). This phenomenon was consistent in all three memory phases (**Figure S18-20**). And the net flow strength of frontal lobe in theta band was larger in match and novel conditions compared to mismatch and old conditions, respectively.

**Figure 9.**
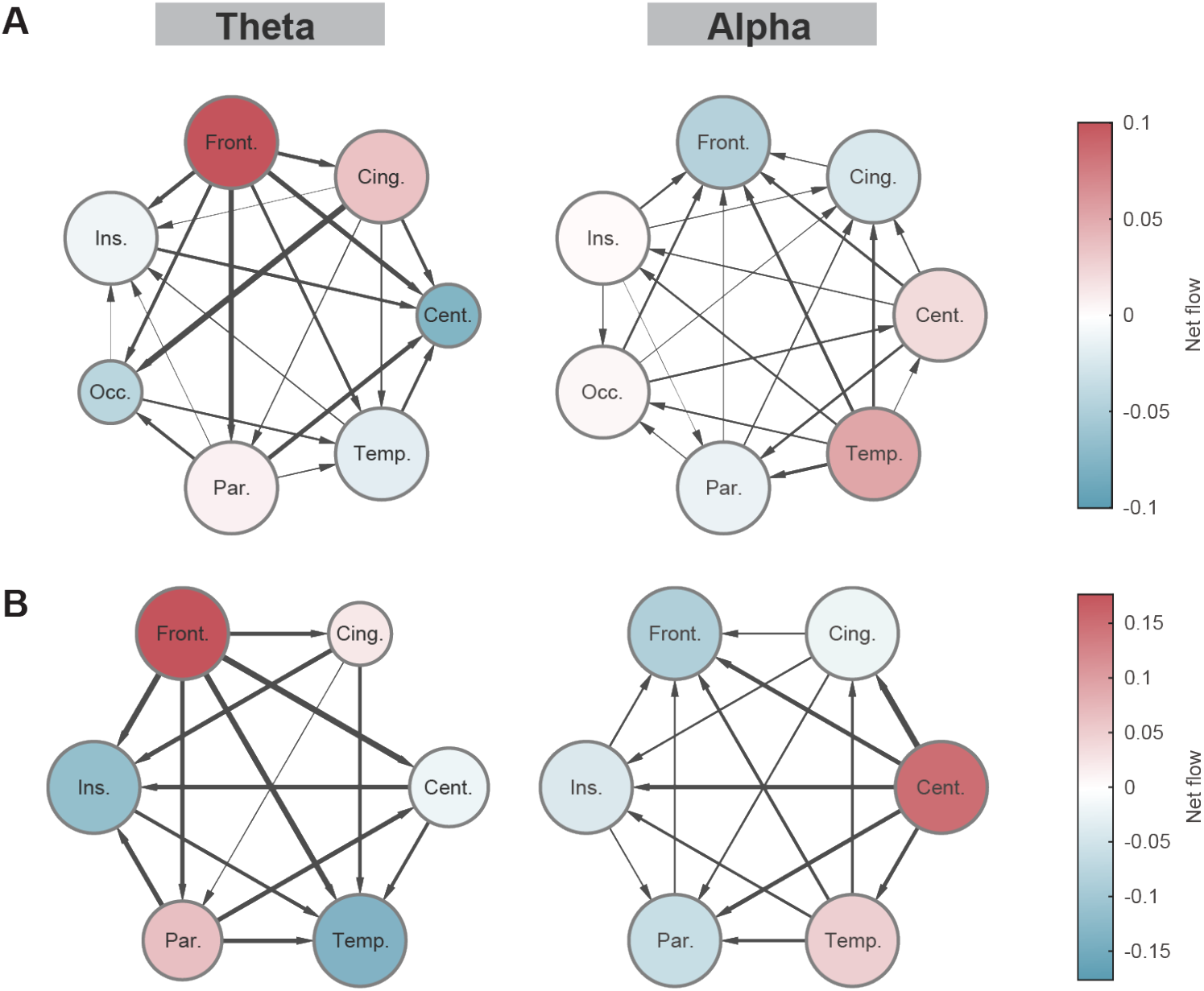
Network in theta and alpha band for all subjects (lobe level, match). Network in theta (first column) and alpha (second column) band for all subjects (**A**) and an example subject (**B**) at lobe level under match condition. Net flow: out-strength minus in-strength. Red circle indicates the node acts as an information source (net flow > 0). Blue circle indicates the node acts as an information sink (net flow < 0). Circle size shows the total connectivity of each node (in-degree + out-degree). The thickness of arrows shows the weights between the nodes.

### TDE-HMM reveals latent task-locked network state trajectories

We modeled task-locked dynamics with a time-delay–embedded hidden Markov model^28,29^ (TDE-HMM, **Methods**). In this framework, each observation at time *t* is represented by a short window of lagged signals, which is then reduced with principal component analysis. The latent states are therefore distinguished by their autocovariance structure rather than by explicit autoregressive parameters, and each state naturally carries a characteristic spectral fingerprint in terms of power and phase coupling. Prior work has shown that this class of HMMs can reliably recover fast, spectrally specific, transient brain states from MEG, EEG, and ECoG data and can be used to compute state-resolved spectra and coherence patterns^30-33^. Guided by this literature, we fixed the number of states to K = 3 for each participant and fit models separately per individual to allow subject-specific state topographies while preserving comparability of state statistics across the group.

For each trial we obtained posterior state probabilities Γ(t, k) and converted them into fractional-occupancy (FO) trajectories by mild temporal smoothing and renormalization. FO summarizes, at each time point, the probability mass carried by each state and has become a standard way to describe HMM dynamics in electrophysiology. The resulting FO time courses (**Figure 10A-B**) revealed rich temporal structure even though the models were blind to task labels during training^31,34^. To capture recurrent patterns of state progression, we aligned trials using dynamic time warping and clustered the FO trajectories via spectral clustering. Across participants this unsupervised procedure consistently returned two robust trajectory motifs (**Figure 10A-B**). The first motif exhibited a pronounced cross-over: one state dominated early in the trial and gradually gave way to a second state in the middle–late portion of the epoch, with a relatively narrow cross-over window around 0.5–0.7 of normalized time. The second motif was comparatively near-stationary, with FO curves remaining flat and no clear exchange of dominance between states. The cross-over motif is reminiscent of stage-like sequences of network states described in previous TDE-HMM work^29,31,34^, whereas the near-stationary motif corresponds to regimes where a single state is occupied for prolonged periods.

**Figure 10.**
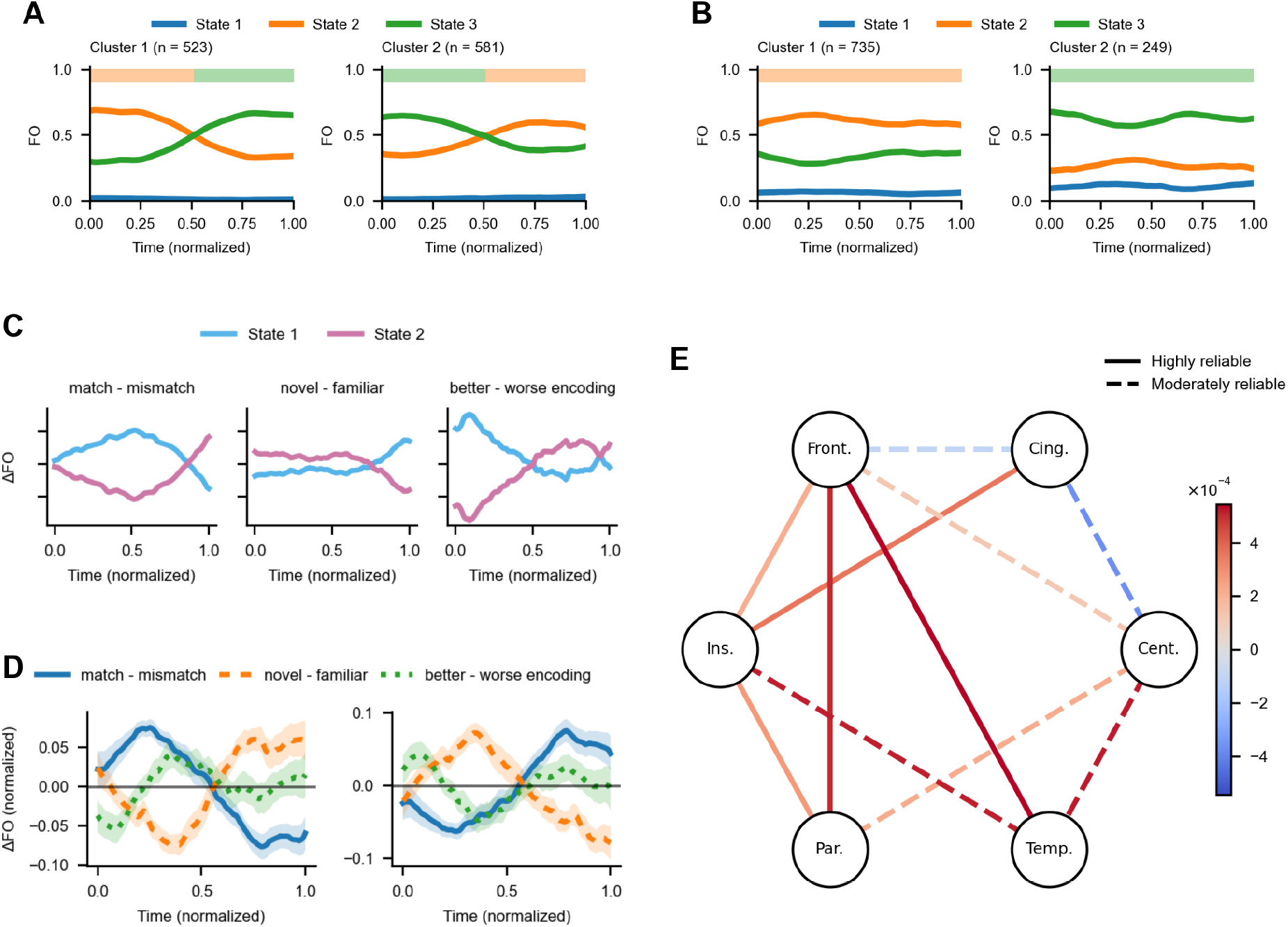
Task-locked HMM states, memory-related trajectories, and lobe-level connectivity. **A-B**. Example fractional-occupancy (FO) time courses of three TDE-HMM states for two participants (A: subject 2; B: subject 6). Subject 2 shows a cross-over motif in which two states exchange dominance once in the middle–late portion of the trial, whereas subject 6 shows a near-stationary motif with relatively flat FO curves and no clear cross-over. **C**. For one example participant, time-resolved differences in FO (ΔFO) between the two memory-related states across three contrasts: retrieval memory (match − mismatch), recognition memory (novel − familiar), and encoding memory (better − worse encoding). **D**. Group-level clustering of ΔFO trajectories from all participants who exhibited either of the two signature motifs. Memory-related states with similar ΔFO profiles are grouped into the same cluster, yielding cross-subject canonical trajectories for retrieval, recognition, and encoding dimensions. Shaded bands indicate SEM across subjects within each cluster. **E**. Lobe-level coherence contrast in the alpha band between the two memory-related network configurations (late-dominant minus early-dominant). Node labels denote cortical lobes; edge color encodes mean Δcoherence, and line style reflects cross-subject reliability (solid: highly reliable; dashed: moderately reliable).

We next examined whether the two memory-related states identified from the FO trajectories were differentially engaged by task conditions. For each participant, we contrasted FO between mismatch and match, novel and familiar, and better and worse encoding trials (ΔFO defined as first minus second condition in each pair) and inspected the time-resolved ΔFO of the two dominant states (**Figure 10C**). Across participants who exhibited the signature trajectories, the two states showed markedly distinct profiles: one state tended to be preferentially expressed in conditions with higher mnemonic or decision demands (e.g., mismatch, novel items, or better-encoding trials), whereas the complementary state was relatively more occupied in the corresponding baseline conditions. These dissociations were robust across all three contrasts and across time within a trial, indicating that the latent states carve up the task into functionally meaningful modes rather than reflecting generic fluctuations in overall excitability.

To align subject-specific states at the group level, we pooled all participants who showed the signature trajectories and clustered the pair of memory-related states from each participant based on their ΔFO time courses across the three contrasts (match − mismatch, novel − familiar, better − worse encoding). This procedure ensured that every participant’s two states were assigned as a pair to one of two canonical families, yielding the group-level prototypes plotted in **Figure 10D**. In both families, the blue curve captures retrieval-memory modulation (match vs. mismatch), the orange curve recognition-memory modulation (novel vs. familiar), and the green curve encoding-memory modulation (better vs. worse encoding). In one family, the retrieval curve shows a pronounced biphasic profile: ΔFO is positive early and becomes negative later in the trial, whereas the recognition curve is approximately anti-phase, starting negative and turning positive in the later period; the encoding curve exhibits a smaller, more centered positive bump. This joint pattern suggests that along the underlying state axis, early portions of the trial favor successful retrieval and familiarity with previously seen items, while later portions increasingly recruit a configuration that is more engaged by novelty and successful encoding, with retrieval-oriented activity relatively suppressed.

In the second family, the three traces retain similar smooth, cross-over shapes but their relative timing and polarity are shifted. Here, the retrieval contrast shows a delayed positive excursion, peaking in the middle–late portion of the trial, whereas the recognition contrast is strongest and positive earlier, then becomes negative toward the response period; the encoding contrast again shows a mid-trial enhancement, but its peak is more tightly aligned with the rise of the retrieval curve. Rather than indicating qualitatively different states, these two families are most parsimoniously interpreted as alternative temporal organizations of the same encoding–retrieval axis: some participants express a trajectory in which recognition- and novelty-related processes are brought online first and are then followed by a retrieval-dominant configuration, whereas others show the converse ordering. This view fits with dynamic-coding accounts in which a limited repertoire of hidden network states is re-used across tasks and subjects, with individual variability in when each state is engaged^35-37^. The coordinated modulation of retrieval (match/mismatch), recognition (novel/familiar), and encoding success (better/worse encoding) along these trajectories is also consistent with ERP and oscillatory work showing partially shared yet temporally staggered mechanisms for novelty detection, recognition decisions, and successful encoding, including mid-frontal and parietal old/new effects and late parietal positivities such as the P3b/LPC^38-41^. Altogether, the clustering results suggest that the TDE-HMM isolates a small set of stereotyped network modes that jointly coordinate retrieval, recognition, and encoding operations, while leaving room for subject-specific shifts in amplitude and timing that likely reflect differences in strategy and task engagement.

Having identified two memory-related network configurations for each participant, we next asked how large-scale connectivity differs between them. Using the HMM state labels, we computed lobe-wise coherence within canonical frequency bands for each configuration and formed a contrast ΔCOH defined as coherence in the late-dominant memory configuration minus coherence in the early-dominant configuration. Positive values therefore indicate stronger coupling when the late configuration is expressed. For each lobe pair and frequency band, we then quantified cross-subject reliability of the sign of ΔCOH: edges were shown with solid lines if there are highly reliable connections and dashed lines indicating moderately reliable connections.

Across bands, a consistent picture emerged (**Figure 10E and Figure S21**). In the beta band, almost all surviving cross-lobar edges were positive, indicating a global tendency for stronger long-range beta coherence in the late-dominant configuration, especially along fronto–parietal, fronto–temporal and insulo–parietal pathways. In the high-gamma band, positive effects were again widespread and centered on temporal cortex, with reliable increases in coherence linking temporal regions to central, parietal and frontal lobes, suggesting that the late configuration engages a more tightly integrated temporo-centric gamma network. Low-gamma coherence showed a more heterogeneous profile: several fronto–parietal and fronto–central edges exhibited positive shifts, whereas insulo–parietal and parietal–temporal connections tended to show reduced coherence in the late configuration, pointing to a possible reweighting rather than uniform strengthening of low-gamma channels. Theta-band effects were sparser and primarily positive, with modest increases along parietal–temporal and insulo– temporal routes. Alpha-band changes were weakest and mixed in sign, with small increases on fronto– parietal and fronto–insular edges but decreased alpha coherence between cingulate and central regions and between parietal and temporal lobes.

This spectral–spatial pattern is broadly consistent with the idea that higher-frequency beta/gamma rhythms support effective long-range communication among task-relevant regions, while slower alpha/theta rhythms provide more selective gating or disengagement of particular pathways^42-45^. In particular, stronger beta and gamma coherence along fronto–temporal and parietal–temporal axes in the late-dominant configuration echoes prior work linking frontoparietal and temporal gamma/beta interactions to controlled memory operations and decision stages^46-48^, while reduced low-gamma or alpha coherence along specific insular and parietal–temporal links may reflect the down-weighting of alternative sensory–context routes once a memory-guided choice is being finalized. Rather than mapping one-to-one onto “encoding” or “retrieval”, these two configurations are thus better understood as complementary large-scale network modes whose relative expression reshapes the routing of information through beta/gamma-mediated channels over the course of a trial.

Together, these analyses show that task-locked neural activity can be parsimoniously described by a small repertoire of latent network states whose temporal progressions, task selectivity, and connectivity profiles are all highly structured rather than random. A TDE-HMM model revealed robust cross-over and near-stationary motifs, within which two memory related states jointly track retrieval, recognition, and encoding demands over time. Clustering across participants aligned these subject specific states into two canonical clusters that differ mainly in the ordering and strength of these memory operations. Finally, lobe-level coherence contrasts demonstrated that transitions between early- and late-dominant configurations are accompanied by frequency specific reweighting of long-range beta/gamma and slower rhythms, suggesting that these latent modes correspond to complementary large-scale network regimes through which memory guided behavior is implemented.

### Integrated summary of functional coupling and network-state analyses

Together, these analyses reveal a distributed but frontotemporal-centered working-memory network whose coupling strength, temporal expression, directional flow, and latent-state organization jointly support recognition, encoding, and retrieval. PAC and PPC first established that condition-dependent FC differences were widespread across the brain. CCI then showed that frontal and temporal regions contributed most consistently to these networks, with parietal and insular regions forming secondary nodes. ERPAC and ERPPC further demonstrated that significant functional connections were temporally structured within trials and showed reproducible cross-subject motifs. dPTE added a directional dimension, indicating frequency-specific source-sink organization across lobes. Finally, TDE-HMM showed that task-locked neural activity could be summarized by a small set of latent network states whose occupancy trajectories and coherence profiles tracked memory operations. This multilevel convergence supports the view that human working memory is implemented through dynamic, directionally organized, large-scale functional coupling networks rather than through isolated regional activation.

## Discussion

Here, we use regionally unbiased analyses to reveal that working memory is implemented through a distributed network. High spatial and temporal resolution intracranial recordings from 36 subjects with 1652 sites allow us to have a more holistic view inside the memory-related manipulating process. We start from capturing the intra- and interareal phase-phase and phase-amplitude couplings, finding that condition-dependent differences are widespread among regions. The frontal and temporal lobes stand out from the coupling structure, implying a key or integrative role during memory processes. The directed networks constructed with phase transfer entropy further illustrate their frequency-specific functions, acting as information hubs. Finally, we depict the latent state trajectories based on hidden Markov model, thereby describing memory-related process at a more schematic level.

Our exploration began with detecting functional couplings among all the recorded sites and multiple frequency bands, for fear that we omitted any uncovered phenomenon happened during the task. Previous studies have made great affords on investigating the individual roles of memory-related regions, such as hippocampus, entorhinal and prefrontal cortex, functioning in memory maintenance, manipulation and computation^49,50^. Region-based studies have been extended to circuit-level as the recording technology advances^51^. Increasingly more research has indicated that functions previously believed to be localized are in fact distributed^52,53^. Within this broader trend, researchers have gradually moved toward explaining the memory process from a network perspective^54-57^. Based on these findings, we tried to ask how these regions serve as the key nodes within the brain-wide memory processes. We therefore designed a more naturalistic paradigm to capture flexible memory behaviors. By reverting to an overview of all the recorded regions, our results support that working memory is a complex process involving the coordinated interactions of multiple regions. This is in line with the view that human association regions are distributed and connected by multiple large-scale networks, which are vital to complex cognitive processes like working memory^12,58,59^.

Unlike sensory-evoked activities which show relatively stable responses in primary sensory regions^60,61^, memory-related activities exhibit more complicated patterns. This complexity manifests in the PAC and PPC results, where condition-dependent differences have been widely found across frequency bands and regions. Even for the well-studied regions such as hippocampus, the highly dynamic patterns make it difficult to explain by simple account. Rather than being interpreted as stimulus representation, power or coupling strength differences between the conditions are more like the observations of different activation patterns. Hence a sole focus on the strength difference leads to a trap under the assumption that one condition should require stronger activity and/or connectivity than another. But naturalistic responses to inputs may employ a large number of the regions for memory processing, which has been confirmed in the first part of our analyses. ERPAC and ERPPC analyses further extended this diversity to the temporal dimension, which evolved with distinct time courses across condition and region pairs. The event-related coupling dynamics suggest that these processes should be unfolded as multidimensional patterns of correlations across regions and time. To further understand the essence of memory, a methodology transition from “strength” to “state” is crucial.

Placing the frontal and temporal lobes back into the memory manipulation network reveals them as prominent nodes, manifested in substantial connectivity and coupling strength with other regions. Their stronger intra-areal correlation fits well with the idea of “small-world”-like organization functional brain networks^62^, which has been extensively explored^63,64^.Frontal and temporal lobes may serve as important nodes within a broader meta-network that supports memory processes^65^. Directed PTE networks further corroborate that the frontal and temporal lobes served as nexuses throughout the task. Information flow diverges from or converges on these regions, differs across frequency bands. The reversed directions in different bands suggest that the same anatomical connection may switch their interaction roles flexibly. Taken together, our results suggest that a complete working memory process is achieved through continuous recurrent interactions across regions, rather than implemented in isolation.

The hidden-state framework provides an advanced way to summarize subject-specific memory dynamics at a more schematic level. We employed TDE-HMM to the signals from all recorded sites without imposing features, location or task labels. The model identified two latent states whose fractional-occupancy trajectories changed along the trial time course in an unsupervised way, offering a description of different memory-related states across the brain. TDE was used for avoiding the inevitable empirical bias derived from manually features selection, as well as allowing spatiotemporal autocorrelation and cross-correlation to contribute to the inferred states. The latent states do not specify which features or correlations drive the observations. They capture the joint structure among the non-independent neural variables. Memory process, from this perspective, can be viewed as transitions among dynamic brain states. Our HMM results provide a complementary state-level support for the coupling analyses and are consistent with state-based accounts of working memory, in which memory processes are maintained and transformed through latent neural states.

Taken together, our results bring us back to a key question in memory field: whether working memory should be considered a temporary storage system, or as a transient computational state. Memory-related periods recruit broad interactions beyond classical regions implicated in memory operations, consistent with state-based accounts of working memory, in which information can be maintained in hidden states and expressed through changes in functional connectivity^35,37,66^. It also holds within the Global Neuronal Workspace Theory (GNWT), where information is first processed within localized processors and becomes globally accessible only if widely broadcasted across the brain^67,68^. In this framework, working memory can be interpreted as a temporary global availability of task-relevant information, so that the activity pattern changes throughout the trials. Flexible memory behavior may depend on the reconfiguration of the connections across regions instead of manipulation within single regions. Our naturalistic paradigm allows us to dissect this flexibility: when recognition, encoding, and retrieval are continuously interleaved, brain organizes memory processing through switching distributed interaction patterns. Future work would like to ask how these states are formed, renewed, and reused across different memory stages.

In conclusion, our analyses converged on a distributed and dynamic view of working memory, in which different memory conditions were expressed through widespread phase-amplitude and phase-phase coupling, phase transfer entropy patterns, and latent state trajectories. Our findings may provide guidance for neural modulation strategies for treating memory disorders in clinical studies. In parallel, these helps constrain biological-inspired computation models based on biological dataset. Our work reshapes how working memory can be understood as an ongoing organization of activity and interaction across the brain, opens a path toward exploring memory process from a dynamic network view, and points to a future in which memory is studied through the flexible coordination of neural systems.

## Methods

### Task paradigm

Participants performed our implementation of a classic card matching game (**Figure 1**), where the objective was to identify all matching pairs by memorizing the position and content of each tile. A board with a dimension of n × n tiles was displayed throughout each block. Initially, all tiles were black and could be clicked to reveal the contents. During each trial, participants clicked and revealed two tiles one after another, at their own pace. Following the second click, if the tiles were of the same image (match), they turned green after 1 second and were no longer clickable for the rest of the current block. If different images (mismatch), the tiles reverted to black/clickable state after 1 second. A block was concluded once all pairs of tiles were matched. The mapping of positions to images remained constant throughout each block. There was no image repetition across blocks. The game commenced on a 3×3 board and escalated to more challenging board dimensions (4×4, 5×5, 6×6, and finally 7×7). Boards with an odd total number of tiles included one distractor object—a human face—without a matching pair. Once reached 7×7, participants remained at this difficulty level for the subsequent blocks. The images used in the task were sourced from the Microsoft COCO 2017 validation dataset^69^ and were displayed in grayscale. The game was programmed and presented using the Psychtoolbox^70,71^ extension in Matlab_R2024b^72^ (Mathworks, Natick, MA) on a 13-inch Apple MacBook Pro laptop.

### Epilepsy patients and recording procedures

Intracranial field potentials were recorded from 36 patients with pharmacologically intractable epilepsy (**Table S2**). Data from 20 patients are publicly available from our previous study^2^. New recordings with the other 16 patients were carried out at Guangdong Sanjiu Brain Hospital (Guangzhou, Guangdong, China) and Shenzhen Second People’s Hospital (Shenzhen, Guangdong, China). Intracranial EEG was recorded using stereo-EEG electrodes and with EEG monitoring systems from Natus (WI, USA) and Nihon Kohden (Tokyo, Japan). All patients were seizure-free during recording sessions. The research protocol was approved by the Institutional Review Boards in both hospitals and Westlake University. The study was conducted with consent from all patients or their legal guardians.

### Behavioral analyses

We calculated some behavioral parameters, such as the reaction time (RT, time between two clicks in a trial), n-since-last-click (the number of clicks since the same tile was clicked), and n-times-seen (number of times the same image had been seen). For n-since-last-click, we excluded trials in which any tile was seen for the first time, i.e., when a tile was not clicked before.

### Electrode localization

We used the iELVis^73^ pipeline for electrode localization. Pre-surgical MR images were segmented using the Freesurfer^74^ software, which were then co-registered with post-implant CT scans. BioImage Suite^75^ was employed to label the electrodes in the MR-CT aligned space. Anatomical locations were automatically returned from FreeSurfer’s brain volume segmentation for depth electrodes. A total of 1,652 electrodes referenced in a bipolar manner were included in the study (**Table S1**). 2,487 electrodes were excluded from analysis due to irrelevant location (white matter, ventricles, pathological sites, etc.), bipolar reference, or having significant noise. The electrode positions were projected onto the MNI305 standard brain model using an affine transformation for visualization purposes.

### Preprocessing of LFP time series

Bipolar reference — a standard referencing method in sEEG studies to minimize volume conduction — was applied to each pair of neighboring recording sites^76^. A zero-phase digital notch filter (Matlab function “filtfilt”) was applied to the bipolarly subtracted LFP signals to remove the AC line noise at 50 Hz (or 60 Hz of data from BCH and BWH) and its harmonics.

### Trial rejection

We identified five categories of questionable trials and excluded them according to the requirements of each analysis. First, we rejected electrode-specific artifact trials based on the trial-wise voltage range. For each electrode, the amplitude range of each trial was calculated as the maximum minus the minimum voltage within the tile-1-aligned analysis window, which extended from 800 ms before the first click to 700 ms after the average first-to-second-click interval. Trials whose voltage range exceeded the across-trial mean by more than 5 standard deviations were marked as artifacts for that electrode. This procedure rejected 0.22% of trials on average across electrodes. Second, we identified abnormal-power trials, defined as trials with unusually large time-frequency power fluctuations. Specifically, for each electrode, normalized power was examined from 5 to 150 Hz after excluding edge time points, and trials whose power range exceeded the across-trial mean by more than 5 standard deviations in any frequency bin were marked as abnormal-power trials. The power spectrum from 5 Hz to 150 Hz (step size = 1 Hz) was computed using the ft_freqanalysis function of the FieldTrip toolbox. We applied the Morlet wavelet transform with 7 cycles per wavelet. The moving window size was frequency-dependent, and the analysis used a time step of 10 ms. These trials accounted for 8.4% of trials on average across electrodes. The union of artifact and abnormal-power trials accounted for 8.6% of trials on average across electrodes.

Third, we excluded out-RT trials, defined as trials with reaction times greater than the subject-level mean plus 5 standard deviations; these accounted for 0.31% of trials on average across subjects. Fourth, we excluded random-match trials from analyses involving associative encoding or retrieval. A random-match trial was defined as a match trial in which the matching partner of the first tile had not been seen previously, such that the correct response could not be guided by associative memory; these trials accounted for 1.09% of trials on average across subjects. Fifth, for retrieval analyses, we excluded excuse-mismatch trials. These were mismatch trials in which the matching partner of the first tile had not been seen previously, and therefore the subject could not have retrieved the correct partner location from memory. The partner-unseen trials identified by this procedure accounted for 30.55% of trials on average across subjects; within the mismatch condition, they were treated as excuse-mismatch trials and excluded from retrieval analyses.

The exclusion criteria were tailored to the memory process being analyzed. Recognition analyses excluded out-RT trials and electrode-specific neural-quality outliers, including artifact and abnormal-power trials. Encoding analyses excluded out-RT trials, random-match trials, and electrode-specific neural-quality outliers. Retrieval analyses excluded all five categories: artifact trials, abnormal-power trials, out-RT trials, random-match trials, and excuse-mismatch trials. In electrode-pair functional-coupling analyses, electrode-specific artifact and abnormal-power trial lists were combined across the two electrodes, and the union of rejected trials was removed before computing coupling measures.

### Working memory processes and conditions

We defined three working memory processes – recognition, encoding, and retrieval, as well as their subordinate conditions by behavioral parameters. Recognition processes were analyzed by separating trials into novel (n-since-last-click = 0) and familiar (otherwise), thereby the novel condition included tiles that were clicked for the first time, and the familiar condition included those that had been revealed before. For WM encoding, we selected only those tiles whose content and location were both novel, to wit, keeping only the 1^st^ flipped tile in each tile pair. The quality of encoding was inferred from n-times-seen-until match (ntsum), and larger ntsum indicated worse encoding. Since subject working memory capacity may vary, a personalized threshold was determined for each individual using unsupervised k-means clustering. For retrieval process, we compared match and mismatch trials, with random match removed. For all functional coupling analysis, only neural signals after the 1^st^ click and before the 2^nd^ click were included.

### Phase-amplitude Coupling

To investigate phase-amplitude coupling^77,78^ (PAC) across brain regions, we extracted low-frequency phase signals and high-frequency amplitude signals from iEEG data. Specifically, theta (5–8 Hz) and alpha (8–14 Hz) bands were used for phase extraction, while low (30–80 Hz) and high (80–150 Hz) frequency bands were used for amplitude extraction. Raw voltage data were band-pass filtered using EEGLAB’s ‘eegfilt’ function for each respective frequency band, followed by Hilbert transform to obtain analytic signals, from which instantaneous phase and amplitude were derived.

For each trial, data were then taken from tile1 to tile2. Disjointed trial segments that pertain to the same condition were linked together to form an unbroken time series for subsequent analyses. The resulting phase and amplitude signals were stored for subsequent PAC analysis.

The Modulation Index (MI) is a metric used to quantify PAC following Tort’s method, where each symbol in its formula has a specific definition: MI refers to the Modulation Index itself, a normalized measure ranging from 0 to 1 that indicates the strength of coupling between low-frequency phase and high-frequency amplitude—with 1 representing maximum coupling and 0 representing no coupling; P_j_ denotes the normalized probability of high-frequency amplitude occurring within the j-th low-frequency phase bin, calculated as the ratio of the mean amplitude in the j-th bin to the sum of mean amplitudes across all bins; A_j_ stands for the mean high-frequency amplitude within the j-th phase bin (phase values here are divided into 18 equal intervals spanning from −180° to 180°, so A_j_ is the average amplitude of the high-frequency signal corresponding to the j-th of these intervals); N is the total number of phase bins, which is fixed at 18 in this study to uniformly segment the full phase range of −180° to 180. The full MI formula is

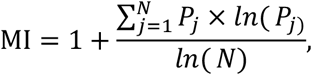

where the probability P_j_ is specifically defined as

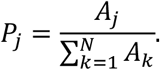

To control for unequal trial counts between two conditions of specific memory phase (e.g., “match” and “mismatch”), the larger set was randomly down-sampled to the size of the smaller set and MI recomputed ten times and averaged; when counts were equal, all data were used. Statistical significance was assessed with a surrogate procedure that circularly shifted the amplitude series relative to the phase, repeated 100 times per electrode pair, frequency combination, and condition while mirroring the same subsampling scheme; one-sided p-values were the proportion of surrogate MIs greater than or equal to the observed MI.

For each electrode pair, condition and frequency combination, we also generated 100 surrogate MI values by circularly shifting the amplitude time series relative to the phase time series, thereby disrupting genuine phase-amplitude alignment while preserving the spectral structure of the signals. The empirical p value was defined as the proportion of surrogate MI values greater than or equal to the observed MI. For each frequency combination, an electrode-pair entry was retained for population-level analysis if the observed MI was significant relative to its surrogate distribution in at least one of the two conditions being compared (empirical p < 0.05).

For population-level analysis, electrodes in specific regions were first mapped to their general regions. Next, for each electrode pair and under each WM condition, the differences in MI of all pairs within the same general region pair were collected and used for Wilcoxon’s signed-rank test to decide whether there was a significant difference between the two conditions. FDR (Benjamini-Hochberg Correction) was applied in all multiple comparisons.

### Phase-phase coupling

We used phase-locking values (PLV) to quantify the strength of phase-phase coupling (PPC) between the five frequency bands (theta: 5–8 Hz, alpha: 8–14 Hz, beta: 14–30 Hz, low gamma: 30–80 Hz, high gamma: 80–150 Hz) across different electrodes. Specifically, we focused on calculating PLV for the same frequency band across distinct electrodes and compared PLV differences under varying experimental conditions.

First, questionable trials were excluded according to the criteria specified earlier (artifact trials, abnormal power trials, out-rt trials, random match trials, and excuse mismatch trials, with specific exclusions tailored to each memory phase analysis). For each remaining trial, we extracted iEEG data spanning from the trial onset to the exact trial offset.

Next, the full dataset (encompassing all retained trials) was band-pass filtered using EEGLAB’s ‘eegfilt’ function for each of the five frequency bands. The Hilbert transform was then applied to the filtered signals to derive instantaneous phase values. Following this, the continuous dataset was segmented into segments corresponding to the two trial types (novel vs. old, better encoding vs. worse encoding, and match vs. mismatch) for each memory phase (recognition, encoding, retrieval). Discontinuous trial segments belonging to the same condition were concatenated to form a continuous time series for subsequent analyses.

We also generated 100 surrogate PLV values for each electrode pair, condition and frequency band by circularly shifting the phase time series of the second electrode relative to the first electrode. This disrupted genuine inter-electrode phase locking. The empirical p-value was defined as the proportion of surrogate PLV values greater than or equal to the observed PLV. For each frequency band, an electrode pair was retained for population-level analysis if its PLV was significant relative to the surrogate distribution in at least one of the two conditions being compared (empirical p < 0.05).

To control for unequal data lengths between conditions, the condition with more samples was randomly subsampled to match the shorter condition, and PLV was averaged across 10 subsampling iterations.

For PLV calculation across electrodes within the same frequency band: The phase difference between signals of the same frequency band from two distinct electrodes was computed. This phase difference was exponentiated by i (the imaginary unit) and summed across all time points, with the magnitude of the sum divided by the total number of time points (N) to yield the PLV^79^:

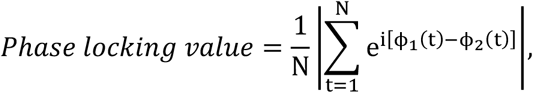

where i is the imaginary unit, ϕ represents the phase in radians, e is the base of the natural logarithm and N is the length of the signal. The range of PLV is [0, 1], where 1 indicates that the phases of the two waves are completely coupled, and 0 indicates no coupling at all.

For population-level analysis, similar to the PAC analysis, electrodes in specific regions were first mapped to their general regions. Next, for each electrode pair and under each WM condition, the differences in PLV of all pairs within the same general region pair were collected and used for Wilcoxon’s signed-rank test to decide whether there was a significant difference between the two conditions. FDR (Benjamini-Hochberg Correction) was applied in all multiple comparisons.

### Connectivity contributive index (CCI)

*CCI* adopts a multi-level scaling procedure which retains the original proportional relationship among regions while accounting for the unbalanced distribution of electrode locations across subjects. For each general region *R*, the *CCI* is calculated as follows. First, for a given frequency combination and a memory process, we computed the ratio of the number of significant region pairs containing *R* to the total number of region pairs containing *R* in the entire dataset, where significant region pairs were those exhibiting significant PAC or PPC difference between memory conditions.

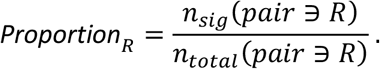

For region pairs where the two regions are the same, it only counts once. Subsequently, we normalized *Proportion*_*R*_ to its largest value in a given frequency combination to yield the *CCI*:

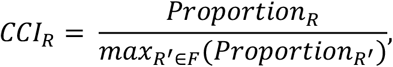

where *F* indicates a given frequency combination. Additionally, if *max* = 0, *CCI* = 0. Lastly, the overall *CCI* of each general region is calculated as the mean value across frequency combinations and then normalized to the largest mean CCI under each memory process.

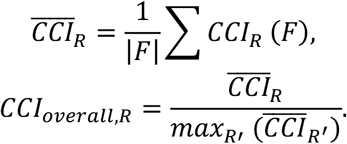

### K-means clustering on ERPAC and ERPPC dynamics

For any region pair previously identified as showing significant differences in PAC (MI, modulation index) or PPC (PLV, phase-locking value) between memory conditions, a surrogate distribution of MI or PLV was generated by calculating 100 times of phase-shuffled MI or PLV. Under each WM process (e.g. recognition), a region pair was excluded from subsequent event-related analysis if p > 0.05 (less than 95 times when the real MI or PLV was larger than the surrogate value) in both conditions (e.g. novel vs. old) of that process. Therefore, the final set of region pairs was an intersection of those that demonstrated significant PAC or PPC difference between conditions and those showing significantly higher MI or PLV than surrogate estimates.

ERPAC (or ERPPC) was computed by a sliding-window MI (or PLV), with a window size of 400 ms and a step size of 25 ms^80^. For each subject, this calculation started from trial onset and ended at the mean reaction time of that subject. ERPAC or ERPPC traces within each group (each panel in **Figure 8-9**) were z-scored to highlight their relative temporal structure. We then applied K-means clustering for time series using dynamic time warping (DTW) as the distance metric (k = 3, 10 iterations, fixed seed) for each group.

### Information flow

For a given frequency band, we applied Hilbert transform to LFP time-series *X*(*t*) to obtain the instantaneous phases *θ*(*t*). Information flow between two electrodes (e.g. *x* and *y*) was inferred from phase transfer entropy (*PTE*)^81^:

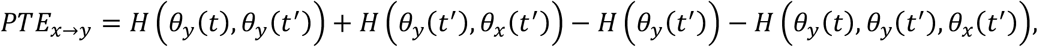

where *x* → *y* indicated the information flow measured with PTE from signal *x* to signal *y. H* stood for Shannon entropy. *θ*_*x*_(*t*’) and *θ*_*y*_(*t*’) were the past phases at time point *t*’ = *t* − *δ*, in which the delay *δ* was calculated following routine procedure^82^:

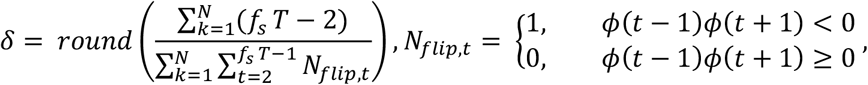

where *f*_*s*_ is the sampling rate (Hz); *T* is the trial duration (s); *N* is the number of electrodes (*N* = 2) in our study. *N*_*flip,t*_ is the number of phase flips between (*t* − 1) and (*t* + 1); phase range was from −*π* to *π*. The unit of *δ* is time point. The marginal entropy and joint entropy were computed as follows:

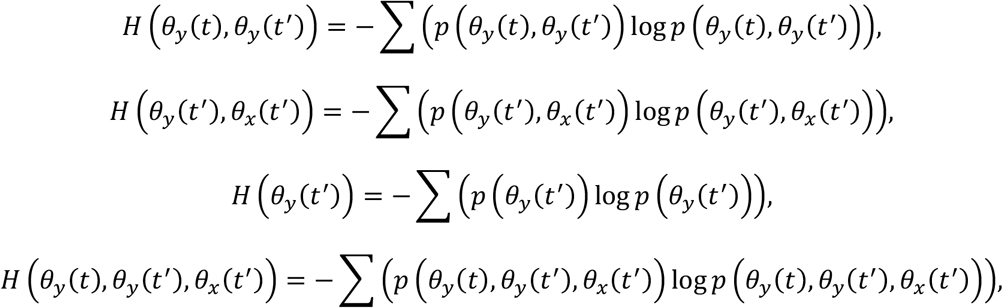

where *H*(*θ*_*y*_(*t*’)) was the marginal entropy for *θ*_*y*_(*t*’), which estimated the uncertainty of the single variable *θ*_*y*_(*t*’). *H*(*θ*_*y*_(*t*), *θ*_*y*_(*t*’)), *H*(*θ*_*y*_(*t*’), *θ*_*x*_(*t*’)), *H*(*θ*_*y*_(*t*), *θ*_*y*_(*t*’), *θ*_*x*_(*t*’)) were the joint entropy, which estimated the uncertainty of multiple variables considered together. *p* represented the probability of the variables. The bin width, or phase resolution, was determined according to previous work^83^:

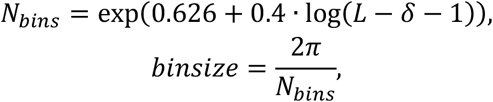

*L* stood for the length of the signal. *PTE* was then normalized to range [0,1] to yield the directed *PTE* (*dPTE*):

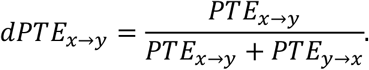

Therefore, the value came to 0.5 if there was no directionality between two signals (symmetric interaction or no significant correlation). We calculated the *dPTE* in delay period (time between the first flip and the second flip) for each trial and for all possible pairs of simultaneously recorded electrodes (i.e., from same subjects). First, Wilcoxon rank-sum test (*H*_0_: the median *dPTE* in two conditions were equal) was performed to determine if there was difference in *dPTE* among different conditions (i.e., novel vs. old, better encoding vs. worse encoding, match vs. mismatch) for each pair of electrodes. Subsequently, for population-level analysis among all subjects, electrodes in specific regions were first mapped to their general regions, for example, orbitofrontal-superiortemporal to frontal-temporal. Next, for each electrode pair and under each WM condition, the median *dPTE* was used as each sample for Wilcoxon signed-rank test to decide whether there was significant difference between conditions in a pair of general regions (*H*_0_: the difference in *dPTE* between two conditions came from a distribution whose median was 0). Lastly, Wilcoxon signed-rank test was performed to determine if *dPTE* exhibited significant directionality (*H*_0_: the *dPTE* under a condition was from a distribution with a median of 0.5). All the tests adopted the Benjamini-Hochberg procedure to control the false discovery rate (FDR) in multiple comparisons.

To pinpoint the information sources and sinks under different conditions and frequency bands, we constructed the directed weighted connectivity matrix to illustrate the connecting strength among anatomical locations. Each node in the matrix represented one location, and each edge symbolled the information flow between two nodes. For a given electrode pair under a given condition, we defined the connectivity weight as “*median dPTE* − 0.5”. Subsequently, the weight between two nodes or regions was computed as the averaged weights of all belonging electrode pairs with significant directionality. The “out-strength” of a given node was the sum of weights from this node to all the other nodes. The “in-strength” of a node was the sum of weights from all the other nodes to this node. The “net flow” of a given node was calculated as its out-strength subtracted by its in-strength. The “in-degree” and the “out-degree” counted the number of edges with significant directionality to and from the node, respectively. General regions recorded in less than 10 subjects were excluded from the analysis.

### Hidden Markov Model

#### Data preparation and standardization

The continuous recordings were first segmented into individual trials. Each trial was then downsampled to reduce computational load and subsequently band-pass filtered between 5 and 120 Hz. After preprocessing, each trial-wise arrays had shape (*n*_channels_ × *n*_samples_). To guarantee valid indexing for every trial, we defined a conservative base dimension as the minimum channel count across non-empty trials and clipped indices accordingly. Trials were resampled to 512hz and concatenated in time to form *X* ∈ *R*^*T*×*N*^ with a trial-boundary array of index-data = [(*t*_0_, *t*_1_), … ], where T denotes the total number of time samples across all concatenated trials and N denotes the number of recording channels. Session-wise standardization followed the HMM-MAR interface.

### Time-delay--embedded HMM (TDE-HMM)^31,84^

#### Observation construction (time-delay embedding)

Let *x*_*i*_ ∈ *R*^*N*^ be a zero-mean multichannel time series (*t* = 1, …, *T*). Choose a symmetric lag set ℒ = {−*τ*_*m*_, …,0, …, *τ*_*m*_}. Form the embedded vector by concatenation:

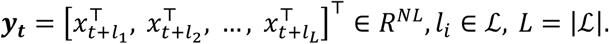

We then apply PCA to {***y***_***t***_} and retain *D* ≪ *N*L components,

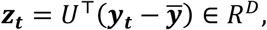

followed by per-component z-scoring, where matrix *U* contains the eigenvectors associated with the *D* largest eigenvalues of the covariance matrix of the embedded vectors. Embedding makes lagged cross-covariances explicit in the observation; as shown below, these carry spectral and phase-coupling information that a Gaussian observation model can encode.

#### Generative model and inference

The latent state sequence {*s*_*t*_} ∈ {1, …, *K*} is a first-order Markov chain with initial distribution π and transition matrix A:

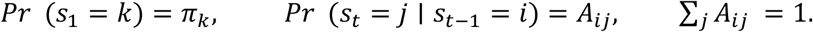

Conditioned on *s*_*t*_ = *k*, the embedded observation Z (in PCA space) is Gaussian with zero mean and (typically diagonal) covariance:

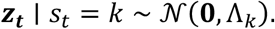

The sequence likelihood is

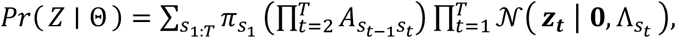

with parameters 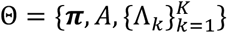. We use the forward-backward algorithm to compute posterior marginals *γ*_*t*_(*k*) = *Pr*( *s*_*t*_ = *k* ∣ *Z*) and pairwise posteriors *ξ*_*t*_(*i, j*) = *Pr*( *s*_*t*−1_ = *i, s*_*t*_ = *j* ∣ *Z*). With diagonal covariances the M-step updates are

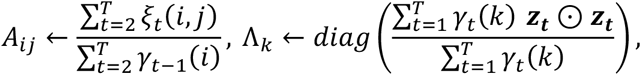

where ⊙ denotes element-wise multiplication, and Λ_*k*_ is the diagonal covariance matrix for state k. We retain Γ(*t, k*) = *γ*_*t*_(*k*) and the Viterbi path for diagnostics. These are the standard EM updates for a Gaussian HMM with diagonal covariances.

#### TDE-HMM is spectrally and phase-coupling aware

Assume that within each state k the process is wide-sense stationary with cross-covariance 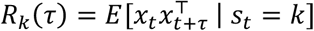. The covariance of ***y***_***t***_, Σ_*k*_ = *Cov*(***y***_***t***_|*s*_*t*_ = *k*),is a block-Toeplitz matrix whose (a, b) block equals *R*_*k*_(l_*a*_ − l_*b*_). By the Wiener-Khinchin theorem, the spectral density is the Fourier transform of *R*_*k*_(*τ*):

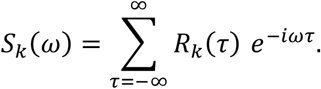

Hence Σ_*k*_ encodes state-specific power and coherence structure; TDE-HMM therefore partitions the data into states with distinct spectral/phase profiles without explicitly fitting multivariate autoregressive model parameters.

#### Implementation choices

We fit the TDE-HMM per participant with K=3 states. Prior to embedding we imputed non-finite values and removed zero-variance features using a threshold of 1e-8. We used symmetric lags covering ±30 ms with stride 2 samples (set by the sampling rate) and applied PCA to retain 90% explained variance; principal components were z-scored. Observation covariances were diagonal with mild Dirichlet/shrinkage prior. Training followed GLHMM APIs^85^; when available, GPU kernels accelerated the forward–backward E-step.

### Trial-wise fractional-occupancy trajectories

Posterior Γ on the embedded axis was mapped back to raw time, and converted to fractional-occupancy (FO) trajectories per trial by applying a centered moving average with a centered 0.5 s window and re-normalizing such that ∑_*k*_ FO(*t, k*) = 1. FO is a standard HMM summary (proportion of probability mass in each state) analyzed alongside switching statistics. Each trial’s FO (t, k) was resampled to a fixed length L= 100 for pairwise comparison.

### Pairwise trajectory dissimilarity using Dynamic Time Warping

To compare FO trajectories by shape while being robust to phase shifts and local stretching, we computed all-pairs distances using DTW. For two trials with multivariate FO trajectories *X* ∈ *R*^*M*×*K*^ and *Y* ∈ *R*^*N*×*K*^, DTW finds a minimal-cost alignment path π and aggregates aligned Euclidean costs:

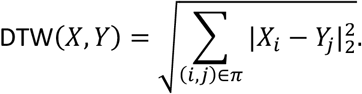

We used a batched implementation with a Sakoe-Chiba band (radius 0.1L) to regularize alignments.

### From distances to a graph and spectral clustering

The DTW distance matrix D was converted to a radial basis function (RBF) affinity

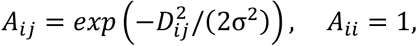

with scale σ set to the median of non-zero upper-triangular distances. We performed spectral clustering following Ng–Jordan–Weiss. Specifically, we formed the normalized Laplacian *L*_sym_ = *I* − *D*^−½^*AD*^−½^, computed the eigenvectors associated with the K smallest eigenvalues, row-normalized the spectral embedding, and ran k-means in the embedded space to obtain trial labels. The number of clusters *K*_*cluster*_ was chosen via the eigengap heuristic within a prespecified range (here, 2–12). Cluster prototypes were visualized as mean *FO*(*t, k*) across member trials.

### Cross-subject clustering of memory-related states

To compare memory-related HMM states across subjects, we first identified for each subject a pair of candidate memory states. These states were chosen based on their ΔFO trajectories and behavioral associations in the single-subject analyses.

For each subject and each state *k* in this pair, we constructed three condition-specific fractional-occupancy (FO) contrasts. Using the smoothed and time-normalized FO trajectories FO(*t, k*)on a common grid of *L* = 100 points, we computed

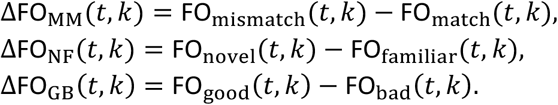

For each contrast and state we subtracted the temporal mean and normalized the resulting curve to unit *ℓ*_2_ norm, so that subsequent clustering was driven by the shape of the ΔFO trajectory rather than its absolute amplitude. The three normalized curves were then concatenated in time to form a 3*L*-dimensional feature vector for that state,

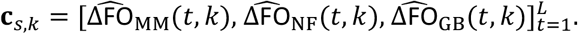

We then clustered these state-specific feature vectors into two families under the constraint that the two states from the same subject must belong to different clusters. To do so, we implemented a custom constrained two-cluster procedure. In each run, we randomly permuted the order of subjects and initialized the two cluster prototypes with the two states of the first subject. We then processed subjects sequentially. For a given subject *s*, we considered the two possible orientations and chose the orientation that minimized the within-cluster squared error to the current prototypes. The prototypes were updated online as running means of all states assigned to each cluster. After all subjects had been assigned, we computed the total within-cluster sum of squared errors. This procedure was repeated for many random orders and both initial orientations (1,000 restarts in the main analyses), and we retained the solution with the lowest total error as the final constrained clustering.

### Lobe-wise coherence analysis

For each subject and HMM state we started from channel-wise coherence spectra *C*(*f*) between all pairs of contacts, as described in the previous section on statewise spectral estimation. Coherence between channels *i* and *j* at frequency *f* was defined as the magnitude-squared coherence

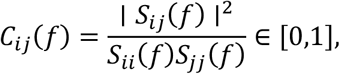

where *S*_*ij*_(*f*)denotes the cross-spectrum and *S*_*ii*_(*f*), *S*_*jj*_(*f*)the autospectra of the two channels.

For a given state, frequency band, and subject, we first averaged the pairwise channel coherence values across all frequency bins within that band, yielding an *N* × *N* coherence matrix *C*^band^(with *N* the number of channels). We then aggregated this matrix at the lobe level. For each pair of lobes (*ℓ*_*i*_, *ℓ*_*j*_), we considered all channel pairs with one contact in *ℓ*_*i*_and one in *ℓ*_*j*_. Cross-lobe coherence was defined as the mean of 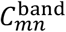 over all such pairs for which channel m and channel n belonged to two different lobes. Within-lobe coherence used only off-diagonal elements (i.e. channel pairs inside the same lobe, excluding self-pairs). This procedure produced, for each state and frequency band, an *t* = *T* × *T* matrix of lobe-wise coherence values, where *T* is the number of lobes. These matrices were exported for statistical analysis and used to generate the lobe-by-lobe coherence heatmaps shown in **Figure 12E** and **Figure S21**.

### Parameter values used in this study

HMM: K=3 states; TDE window ±30 ms (stride 2); PCA 90%; diagonal covariances with mild prior; per-subject training; GPU-accelerated E-step when available. FO extraction: 0.5 s centered smoothing; unit-sum normalization; resampling to L=100. DTW: batched cross-similarity; Sakoe–Chiba radius = 0.1L; optional soft-DTW.

Spectral clustering: RBF affinity with *σ* = median{*D*_*ij*_: *i* < *j, D*_*ij*_ > 0};*L*_sym_; Ng–Jordan–Weiss; K by eigengap (2–12).

## Acknowledgement

We thank Jiemin Jia for insightful discussions and feedback on the project; Xinjing Li and Hao Zhu for guiding equipment setup; Xing Tian for sharing equipment; and all patients and their families for their generosity to support this study. The research was funded by the Westlake Fellows Program at Westlake University and Funding No. JCYJ20240813141105007 supported by Shenzhen Science and Technology Innovation Committee, China.

## Funder Information Declared

Westlake University, https://ror.org/05hfa4n20, Westlake Fellow Program

