## Supplemental Tables for "Large-scale functional coupling and computational modelling reveal frontotemporal hotspots in distributed network connectivity during working memory recognition, encoding, and retrieval"

**Table S1**

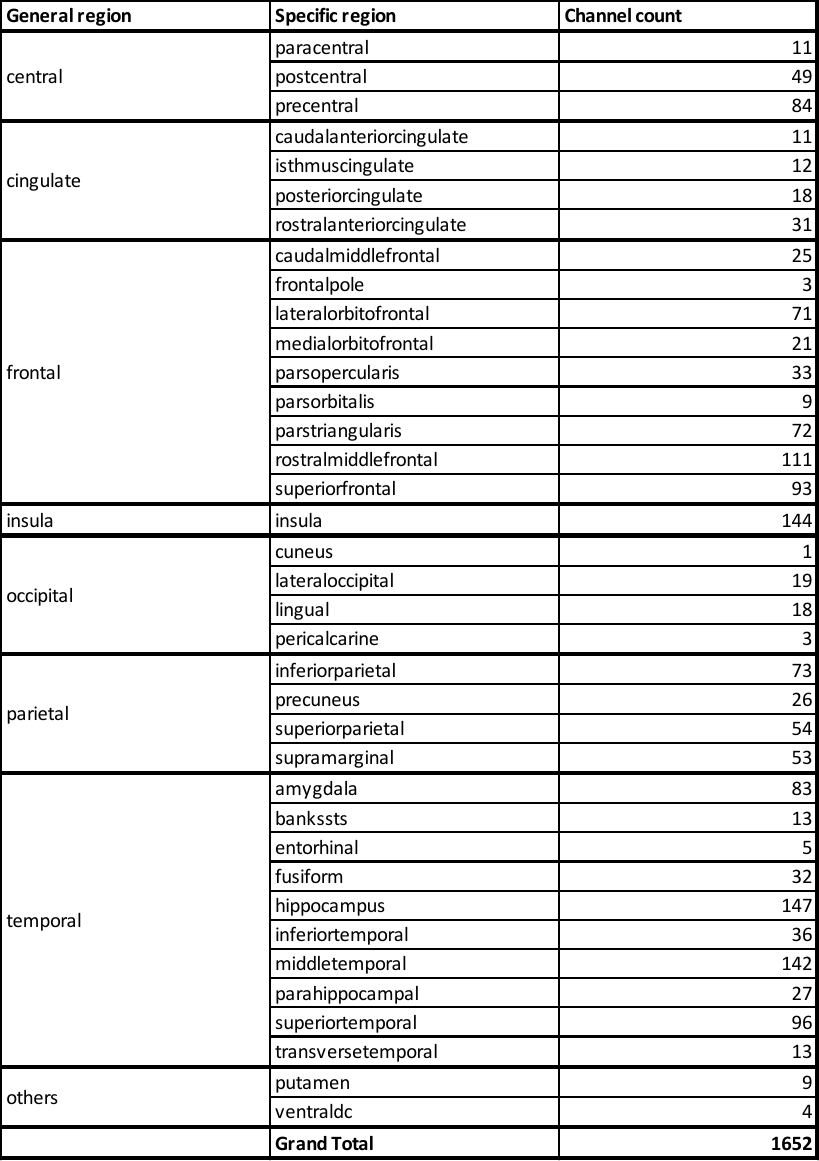

**Table S1. Anatomical locations of all channels recorded.**

**Table S2**

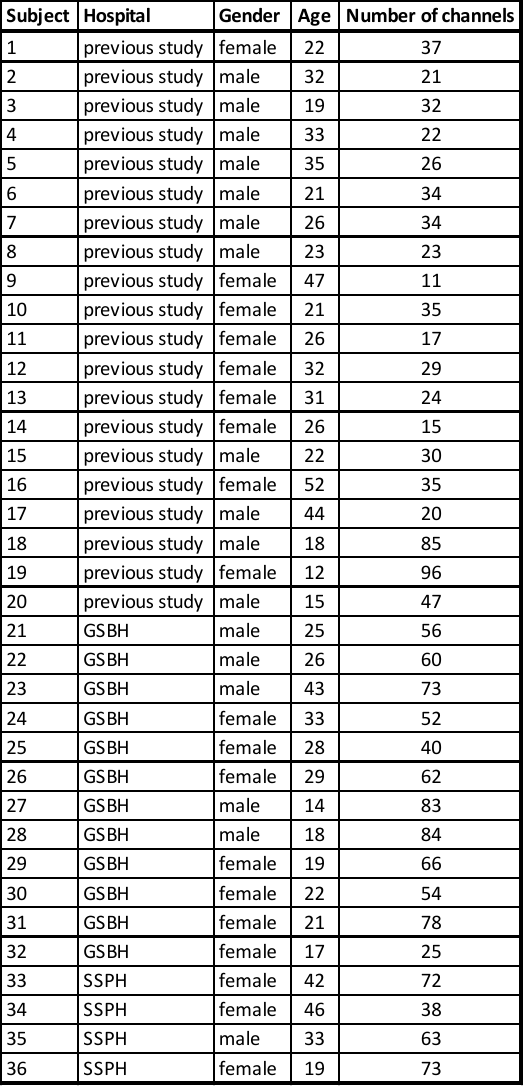

**Table S2. Subject information.** Both electrophysiological data and anatomical information of subjects 1-20 are publicly available at <https://klab.tch.harvard.edu/resources/HowToGetAMatch.html>. Data of other subjects were collected at Guangdong Sanjiu Brain Hospital (GSBH, Guangzhou, Guangdong, China) and Shenzhen Second People’s Hospital (SSPH, Shenzhen, Guangdong, China).
