## Supplemental Figures for "Large-scale functional coupling and computational modelling reveal frontotemporal hotspots in distributed network connectivity during working memory recognition, encoding, and retrieval"

### Supplementary: Figure S1

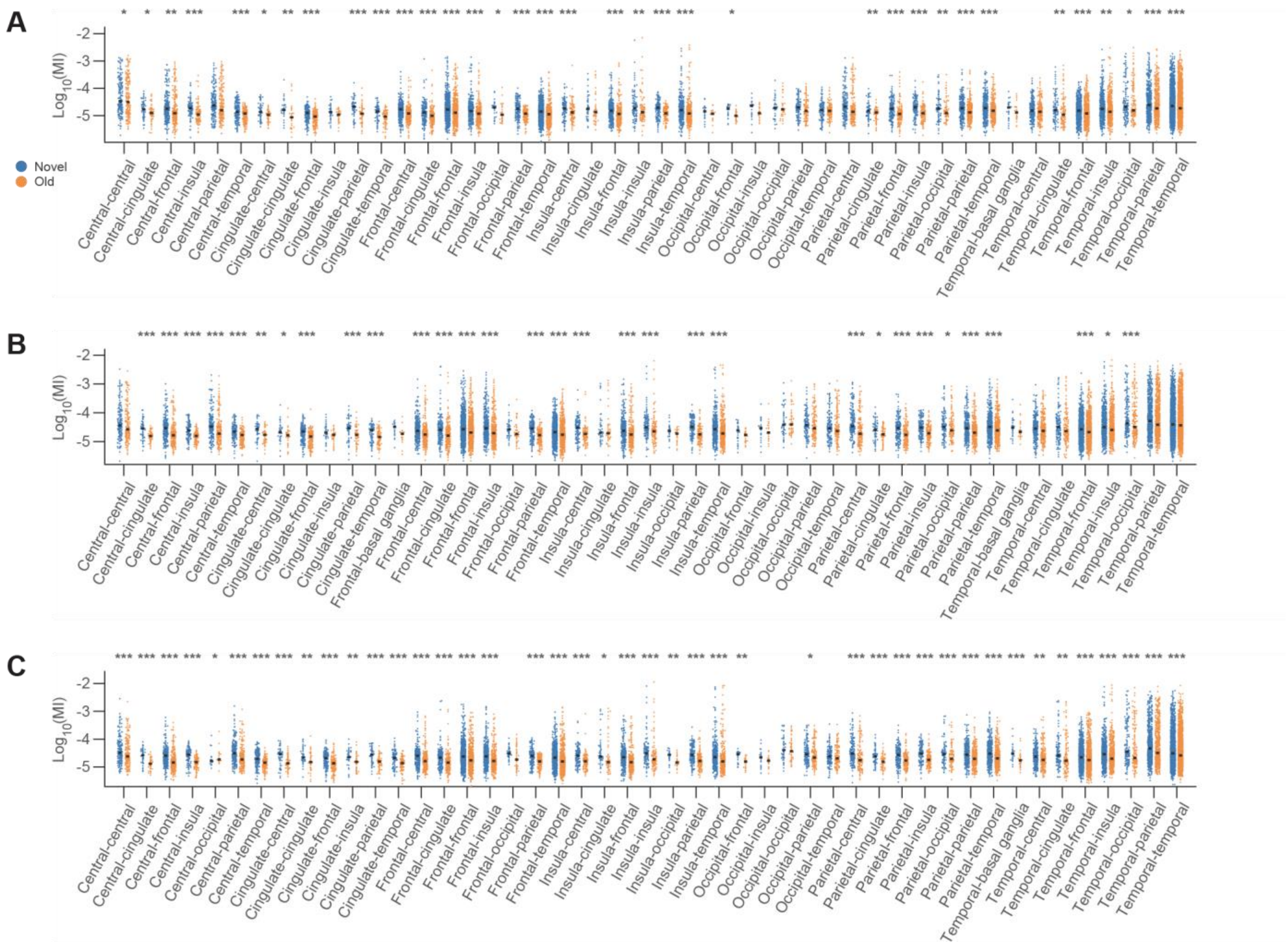


**Figure S1. Widely distributed phase-amplitude coupling under recognition process. (A)** Alpha-high gamma, **(B)** theta-low gamma and **(C)** theta-high gamma PAC strength in each combination of general regions from all subjects during recognition. Modulation index (MI) is shown in log10 scale to account for its exponential distribution. Black horizontal bars indicate the median PAC value in each condition. Location combinations containing less than 20 samples were removed from analysis. The asterisks highlight region pairs with statistically significant differences between conditions (Wilcoxon signed-rank test, $\alpha = 0.05$, *$p < 0.05$, **$p < 0.01$, and ***$p < 0.001$. All p-values were corrected for multiple comparisons using the false discovery rate (FDR) method.

### Supplementary: Figure S2

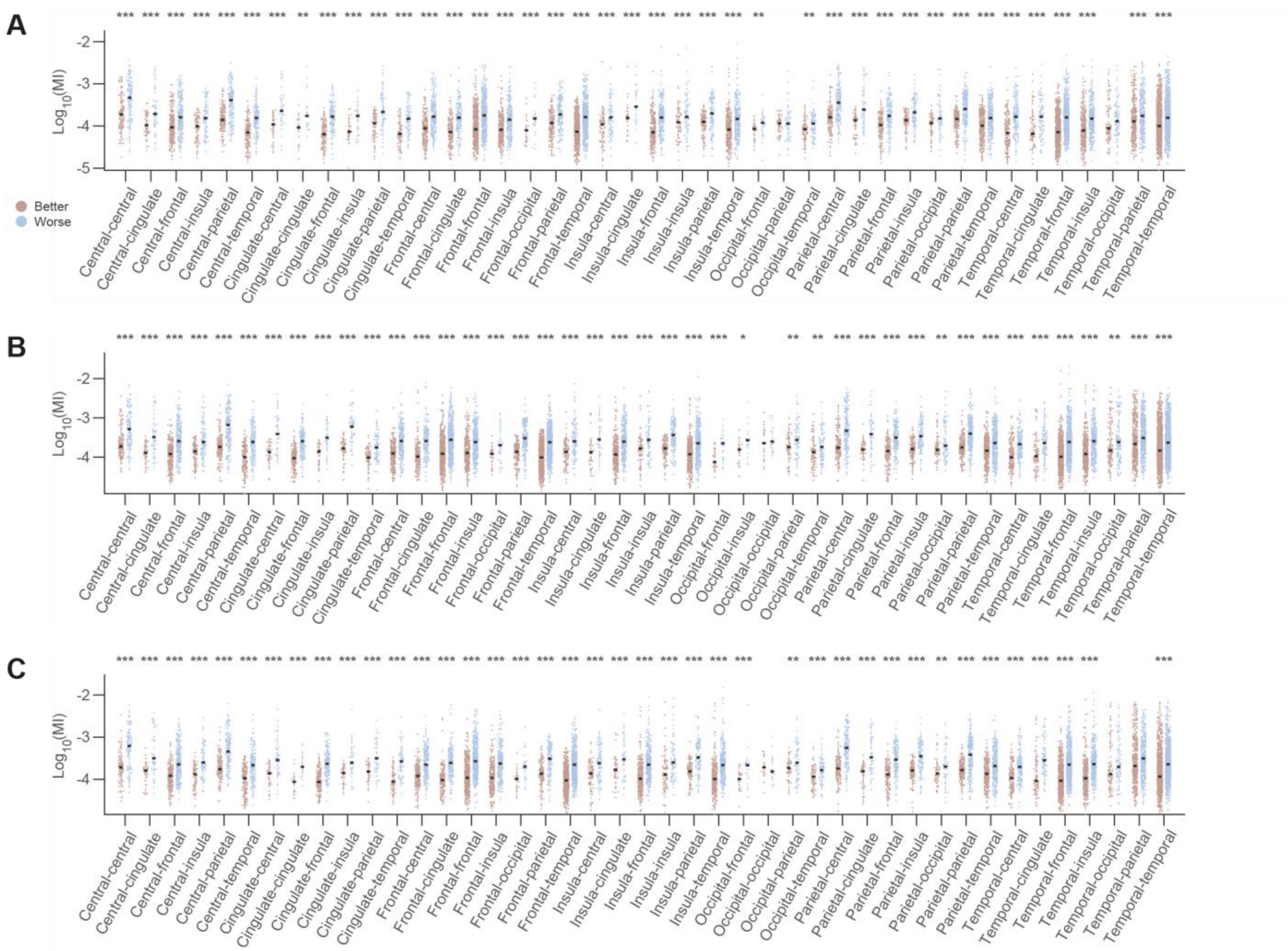


**Figure S2. Widely distributed phase-amplitude coupling under encoding process. (A)** Alpha-high gamma, **(B)** theta-low gamma and **(C)** theta-high gamma PAC strength in each combination of general regions from all subjects during encoding. For figure formats, follow figure S.

### Supplementary: Figure S3

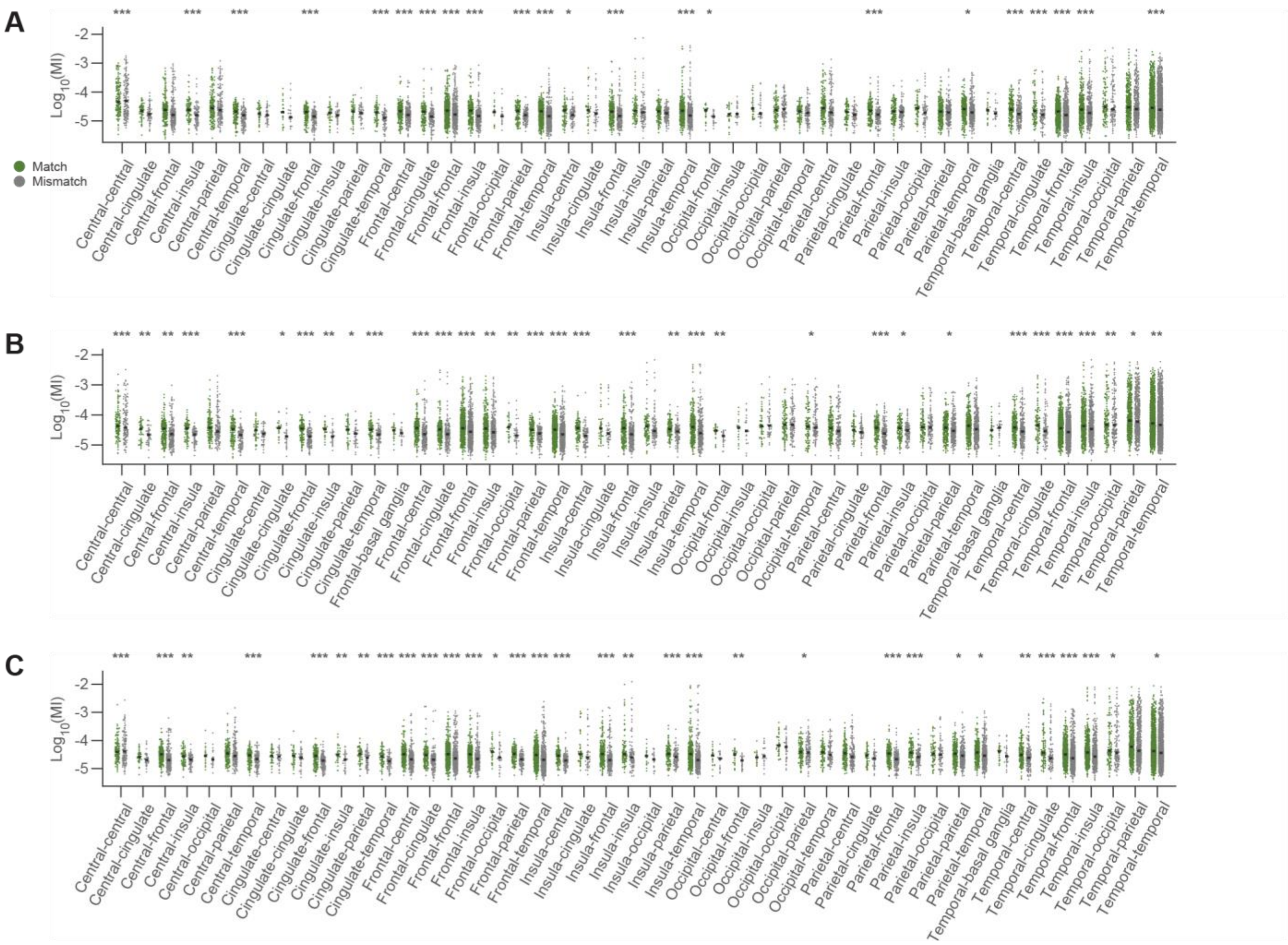


**Figure S3. Widely distributed phase-amplitude coupling under retrieval process.** **(A)** Alpha-high gamma, **(B)** theta-low gamma and **(C)** theta-high gamma PAC strength in each combination of general regions from all subjects during retrieval. For figure formats, follow figure S.

### Supplementary: Figure S4

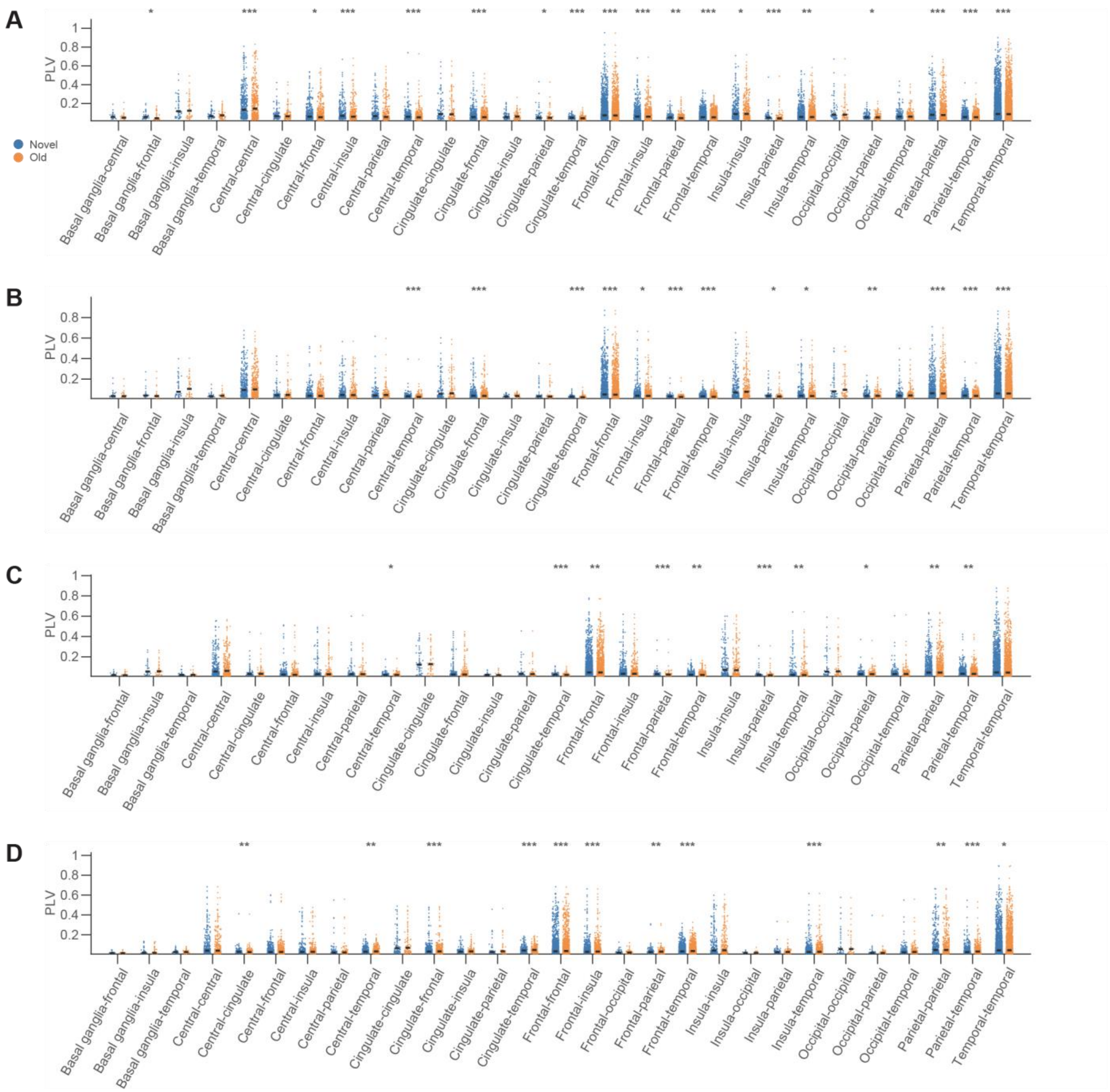


**Figure S4. Widely distributed phase-phase coupling under recognition process. (A)** Alpha, **(B)** beta, **(C)** low gamma and (**D**) high gamma PPC strength in each combination of general regions from all subjects during recognition. Black horizontal bars indicate the median PPC value in each condition. Location combinations containing less than 20 samples were removed from analysis. The asterisks highlight region pairs with statistically significant differences between conditions (Wilcoxon signed-rank test, $\alpha = 0.05$, *$p < 0.05$, **$p < 0.01$, and ***$p < 0.001$. All p-values were corrected for multiple comparisons using the false discovery rate (FDR) method.

### Supplementary: Figure S5

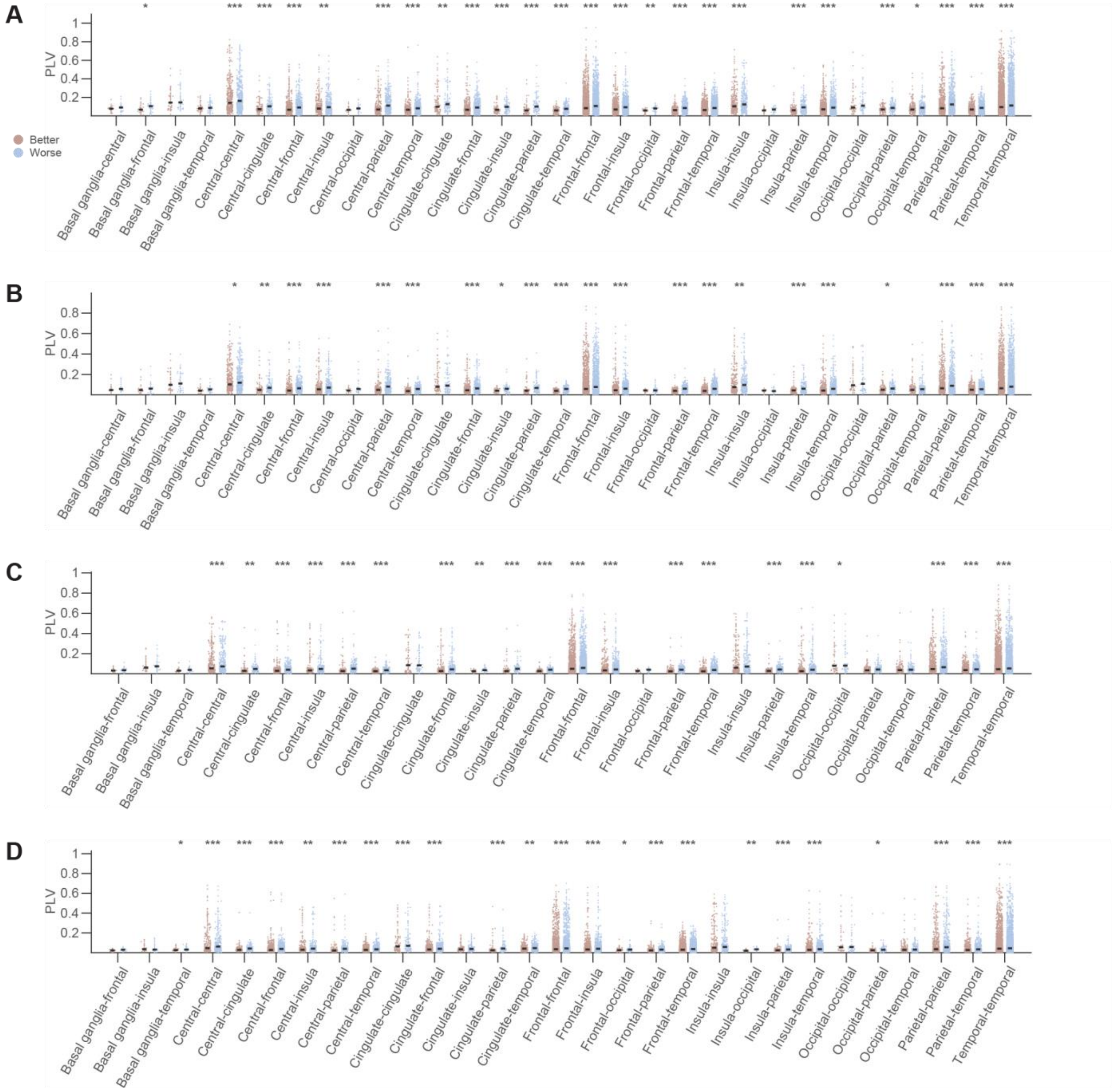


**Figure S5. Widely distributed phase-phase coupling under encoding process. (A)** Alpha, **(B)** beta, **(C)** low gamma and (**D**) high gamma PPC strength in each combination of general regions from all subjects during encoding. For figure formats, follow figure S.

### Supplementary: Figure S6

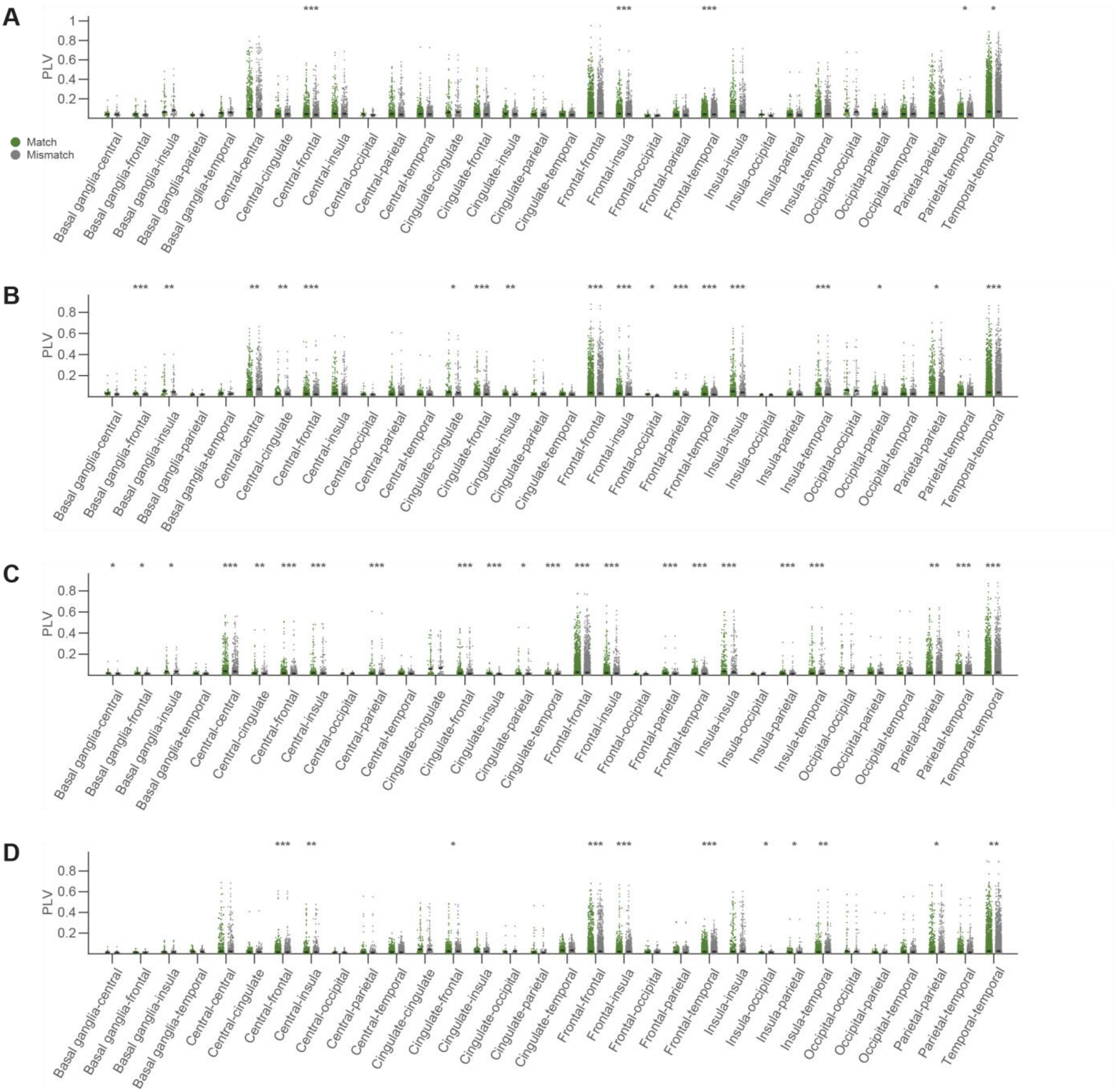

**Figure S6. Widely distributed phase-phase coupling under retrieval process. (A)** Alpha, **(B)** beta, **(C)** low gamma and (**D**) high gamma PPC strength in each combination of general regions from all subjects during retrieval. For figure formats, follow figure S.

### Supplementary: Figure S7

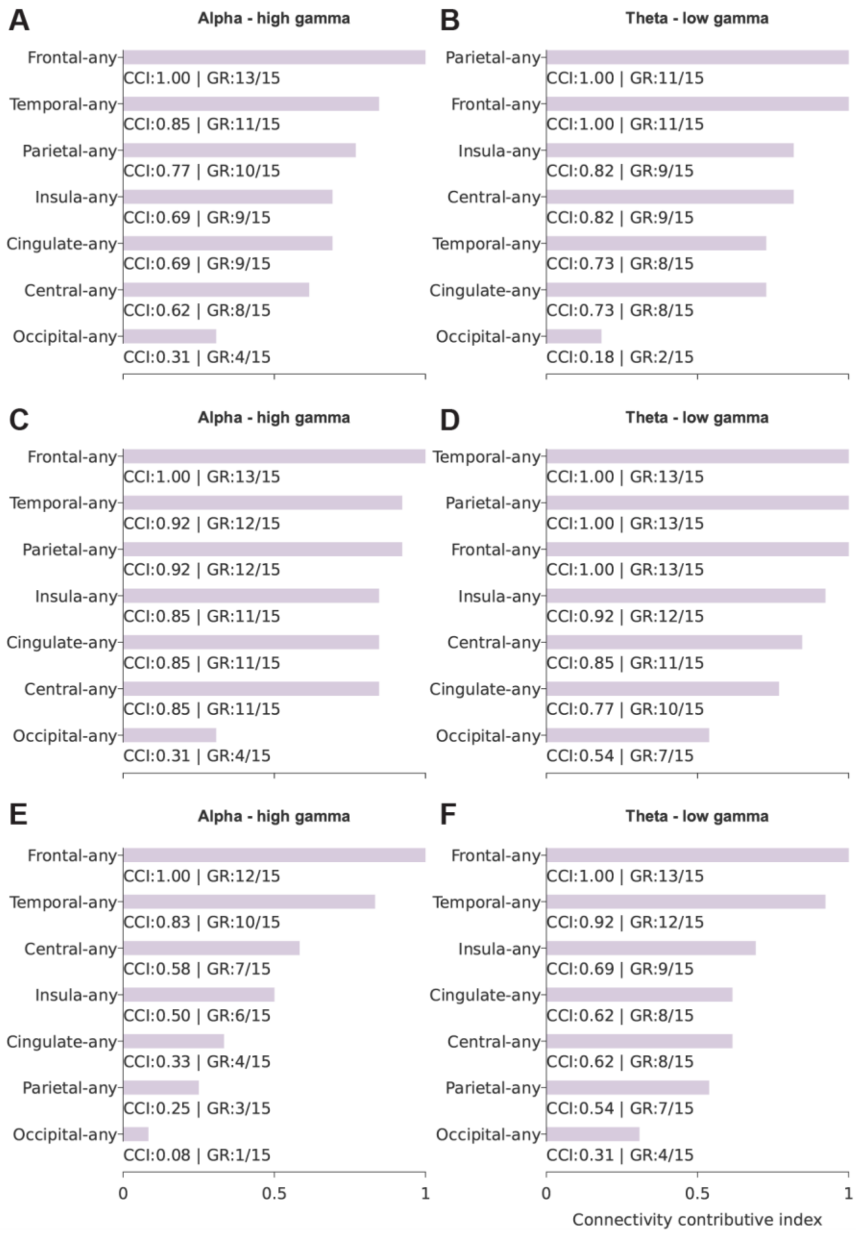


**Figure S7. Regional contribution to PAC network.** The contribution of any general region to the overall PAC network was estimated using connectivity contributive index (CCI). Each row shows CCI results of each WM process. **A-B**: recognition; **C-D**: encoding; **E-F**: retrieval. The left column displays results in alpha-high gamma PAC and the right column theta - low gamma PAC. Figure formats follow those in **Figure 4**.

### Supplementary: Figure S8

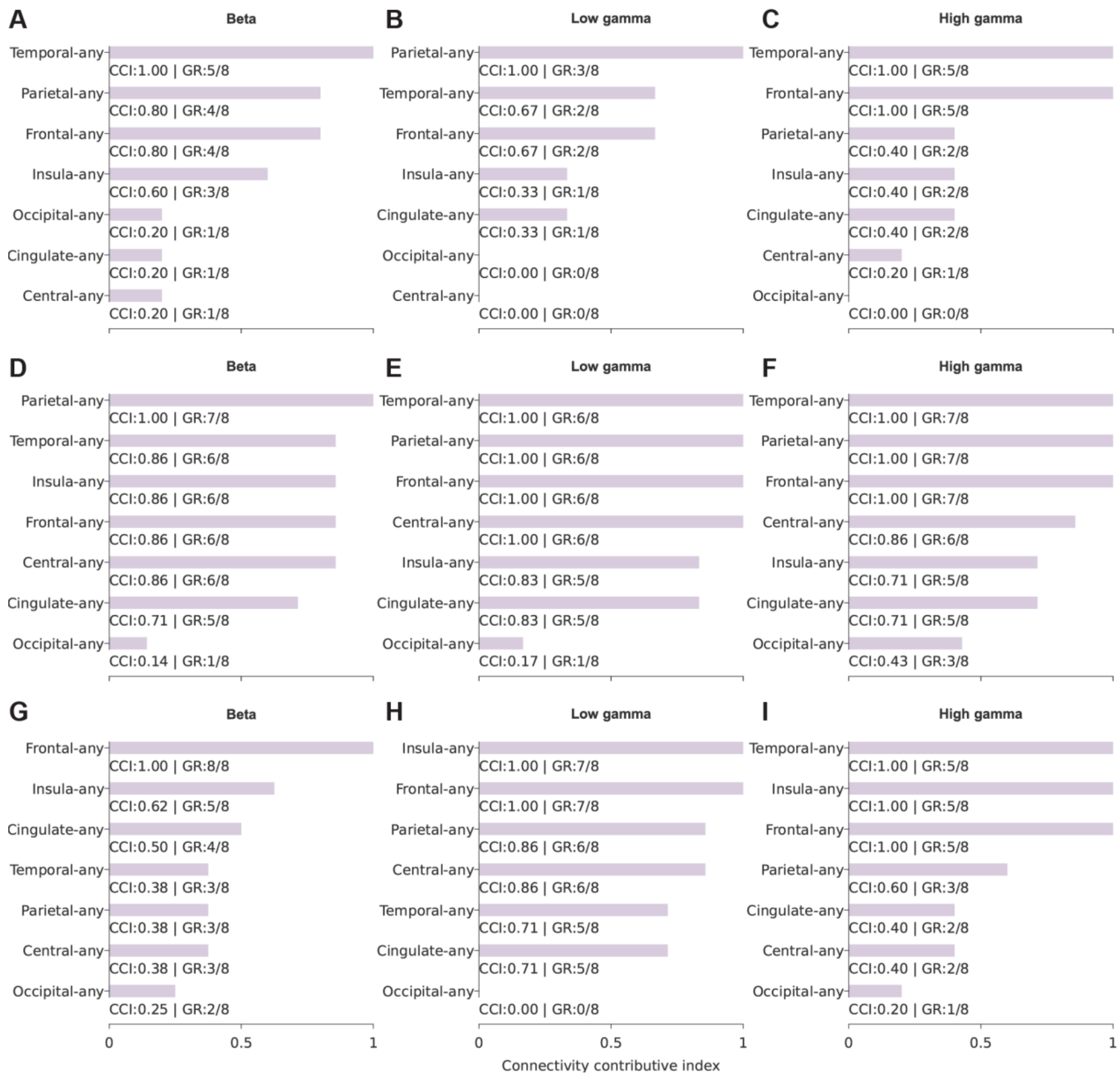


**Figure S8. Regional contribution to PPC network.** The contribution of any general region to the overall PPC network was estimated using connectivity contributive index (CCI). Each row shows CCI results of each WM process. **A-C**: recognition; **D-F**: encoding; **G-I**: retrieval. The Left column displays results in beta PPC, the middle column low gamma PPC, and the right column high gamma PPC. Figure formats follow those in **Figure 5**.

### Supplementary: Figure S9

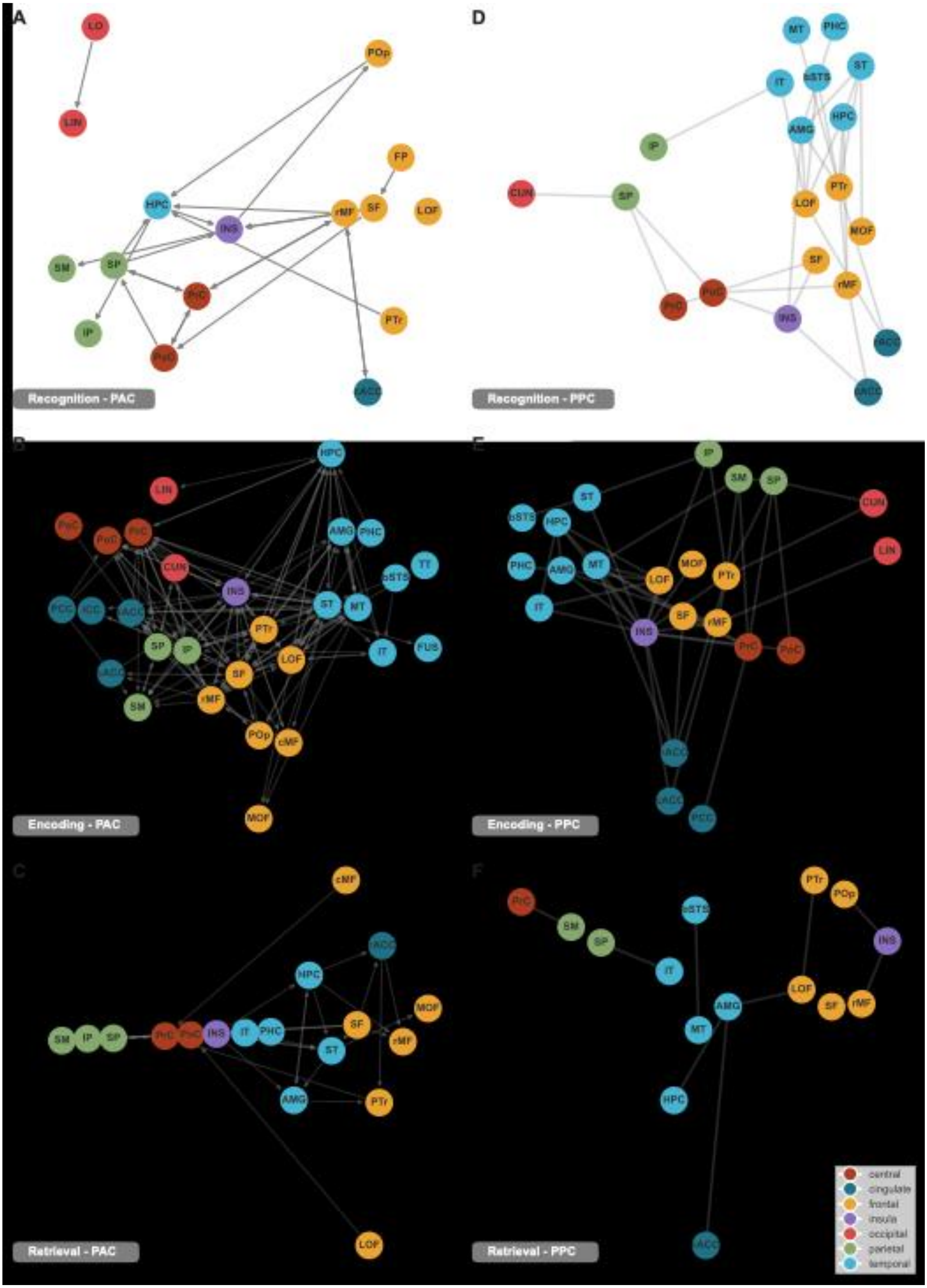

### Figure S9 Caption

**Figure S9. Overview of PAC and PPC networks during recognition, encoding, and retrieval. A-C.** Brain regions showing alpha – low gamma PAC difference (Wilcoxon signed rank test, FDR corrected $p < 0.05$) between novel and old (**A**), better and worse encoding (**B**), as well as match and mismatch (**C**). Each node represents one specific brain region, and the color indicates the general region it belongs to. Edges connect region pairs that exhibited PAC differences. Arrows label the coupling direction, where the origins are the regions for phase, and the destinations regions for amplitude. **D-F**. Brain regions theta PPC difference (Wilcoxon signed rank test, FDR corrected $p < 0.05$) between novel and old (**D**), better and worse encoding (**E**), as well as match and mismatch (**F**). Edges connect region pairs that exhibited PPC differences. PaC: paracentral. PoC: postcentral. PrC: precentral. cACC: caudalanteriorcingulate. ICC: isthmuscingulate. PCC: posteriorcingulate. rACC: rostralanteriorcingulate. cMF: caudalmiddlefrontal. FP: frontalpole. LOF: lateralorbitofrontal. MOF: medialorbitofrontal. POp: parsopercularis. POr: parsorbitalis. PTr: parstriangularis. rMF: rostralmiddlefrontal. SF: superiorfrontal. INS: insula. CUN: cuneus. LO: lateraloccipital. LIN: lingual. Peri: pericalcarine. IP: inferiorparietal. PCU: precuneus. SP: superiorparietal. SM: supramarginal. AMG: amygdala. bSTS: bankssts. EC: entorhinal. FUS: fusiform. HPC: hippocampus. IT: inferiortemporal. MT: middletemporal. PHC: parahippocampal. ST: superiortemporal. TT: transversetemporal.

### Supplementary: Figure S10

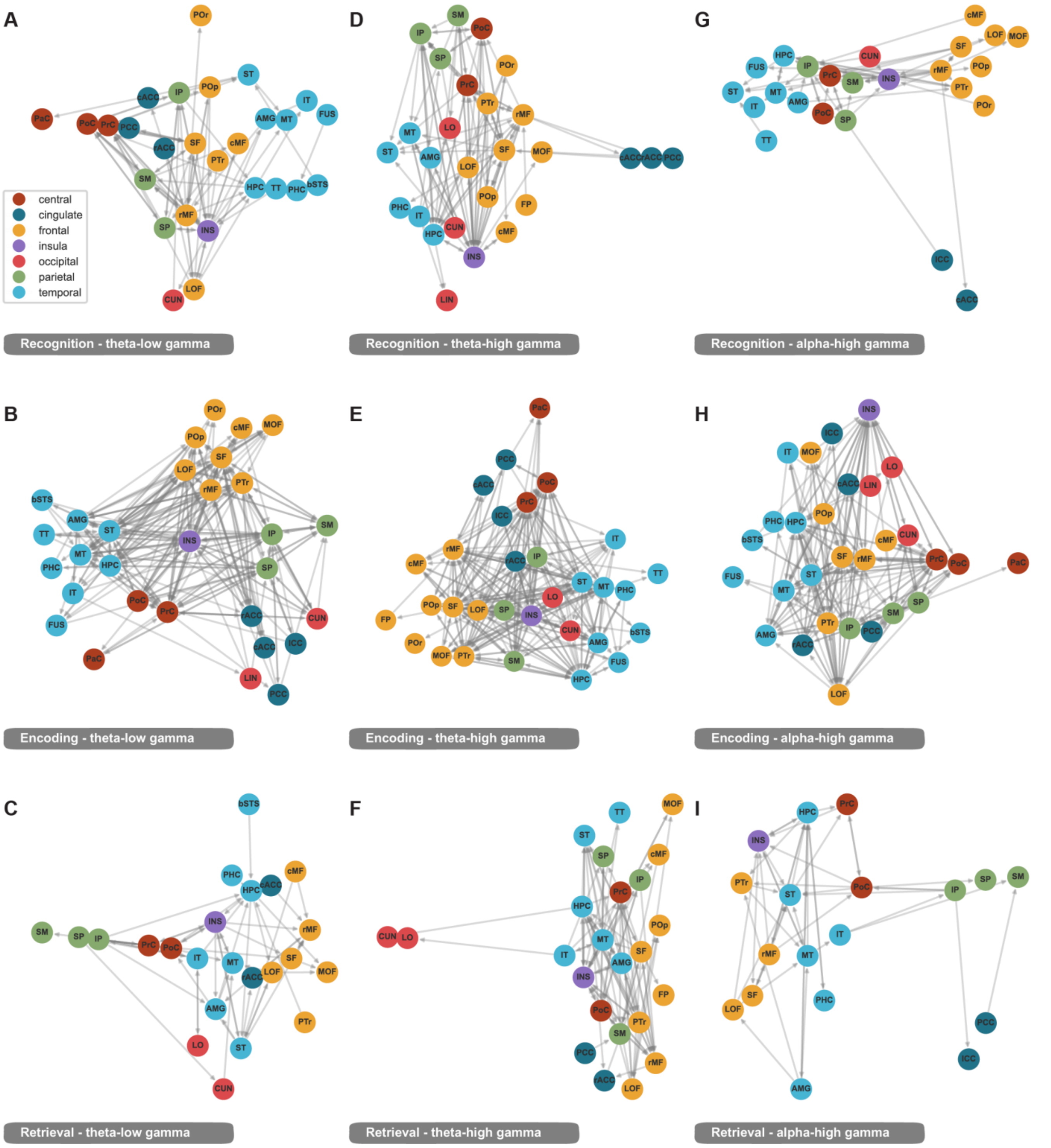

### Figure S10 Caption

**Figure S10. Overview of PAC networks during recognition, encoding, and retrieval. A-C.** Brain regions showing theta – low gamma PAC difference (Wilcoxon signed rank test, FDR corrected $p < 0.05$) between novel and old (**A**), better and worse encoding (**B**), as well as match and mismatch (**C**). **D-F**. Brain regions theta – high gamma PAC difference (Wilcoxon signed rank test, FDR corrected $p < 0.05$) between novel and old (**D**), better and worse encoding (**E**), as well as match and mismatch (**F**). **G-I**. Brain regions alpha – high gamma PAC difference (Wilcoxon signed rank test, FDR corrected $p < 0.05$) between novel and old (**G**), better and worse encoding (**H**), as well as match and mismatch (**I**). Figure formats follow those in Figure 6.

### Supplementary: Figure S11

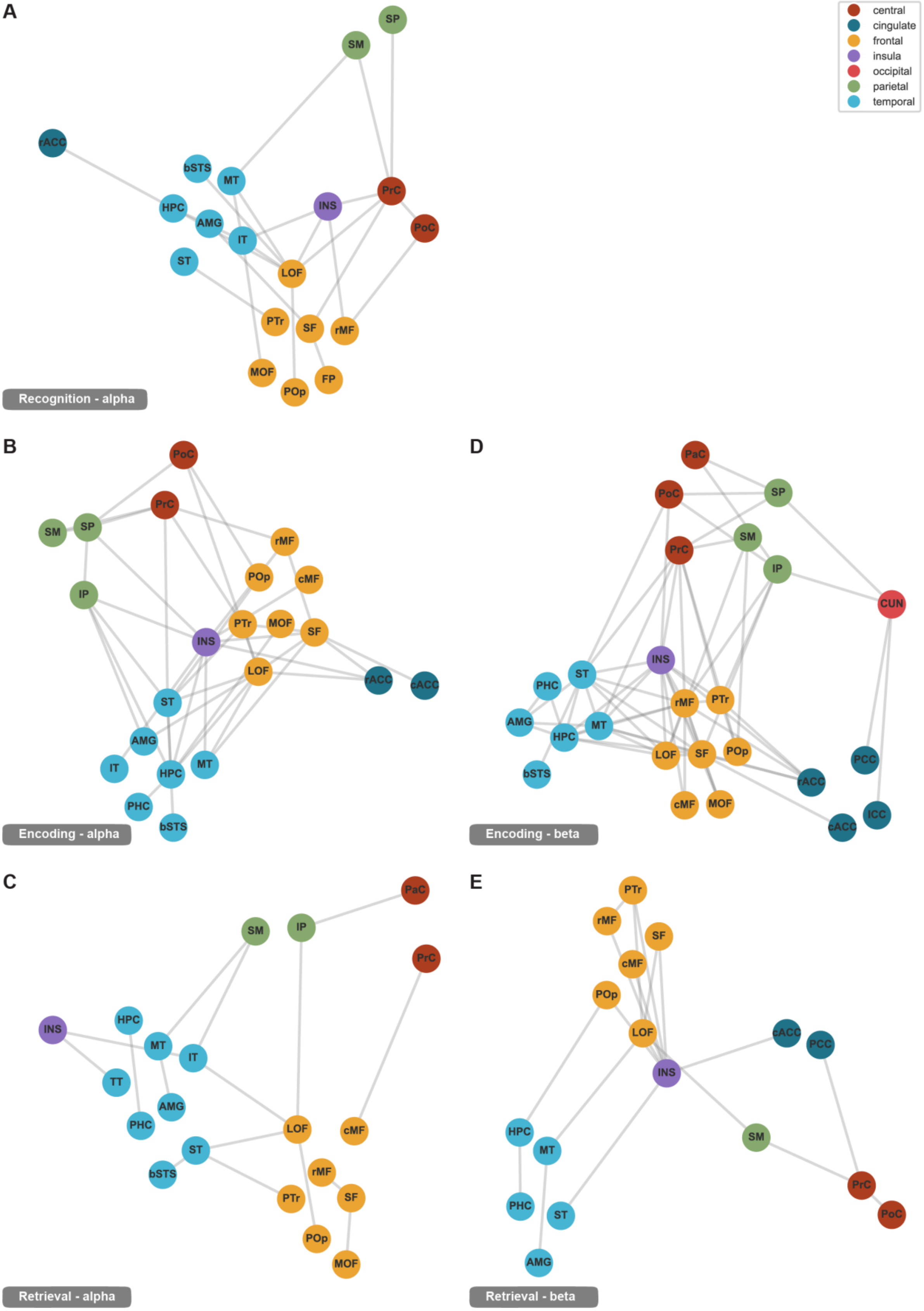

### Figure S11 Caption

**Figure S11. Overview of PPC networks in low frequency bands. A-C.** Brain regions showing alpha PPC difference (Wilcoxon signed rank test, FDR corrected $p < 0.05$) between novel and old (**A**), better and worse encoding (**B**), as well as match and mismatch (**C**). **D-E**. Brain regions beta PPC difference (Wilcoxon signed rank test, FDR corrected $p < 0.05$) between better and worse encoding (**D**) as well as match and mismatch (**E**). No region pair showed significant beta PPC difference between novel and old. Figure formats follow those in Figure 6.

### Supplementary: Figure S12

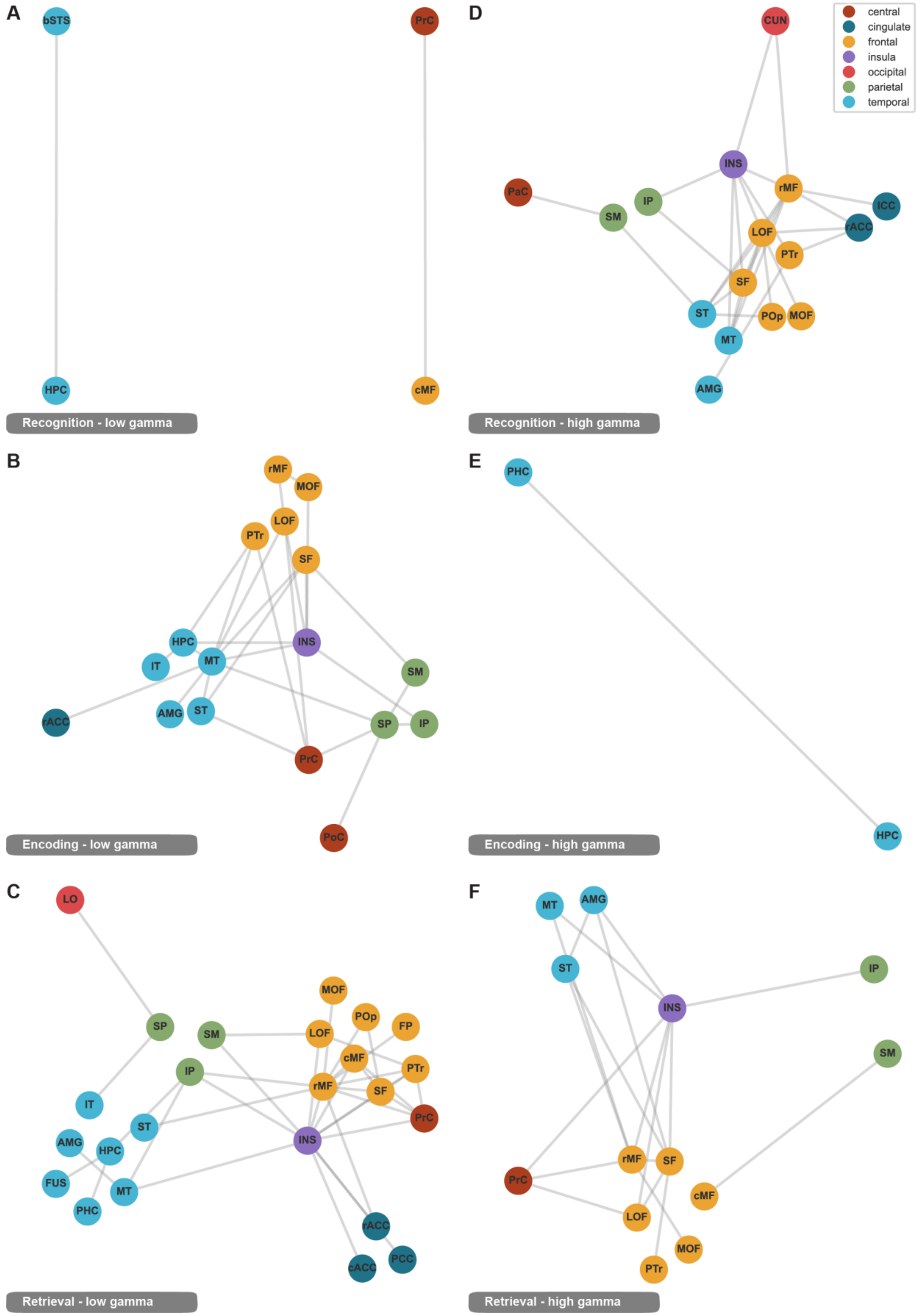

### Figure S12 Caption

**Figure S12. Overview of PPC networks in high frequency bands. A-C.** Brain regions showing low gamma PPC difference (Wilcoxon signed rank test, FDR corrected $p < 0.05$) between novel and old (**A**), better and worse encoding (**B**), as well as match and mismatch (**C**). **D-F**. Brain regions high gamma PPC difference (Wilcoxon signed rank test, FDR corrected $p < 0.05$) between novel and old (**D**), better and worse encoding (**E**), as well as match and mismatch (**F**). Figure formats follow those in Figure 6.

### Figure S13

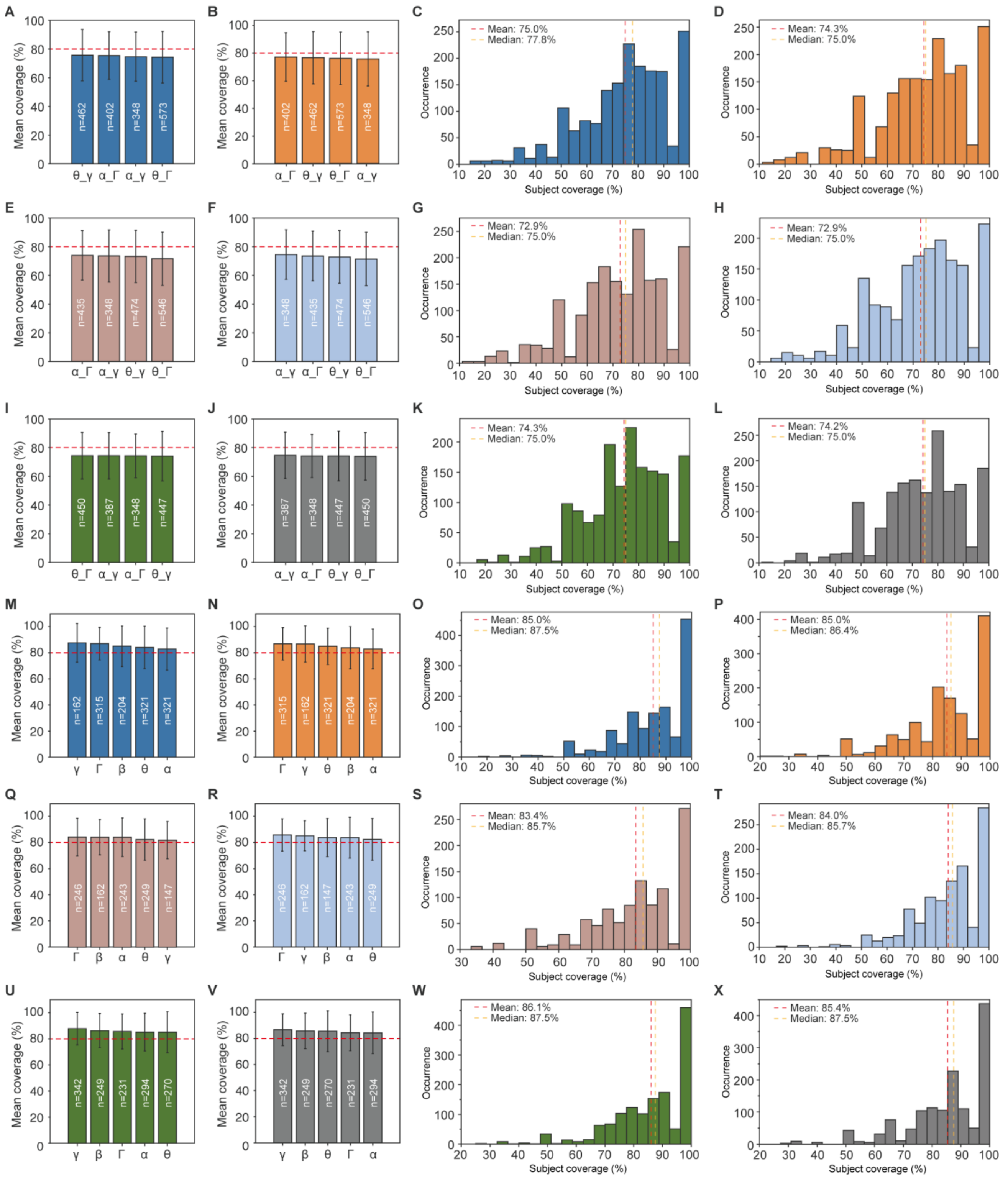

A
Mean coverage (%)
n=462
n=402
n=348
n=573
θ_γ α_Γ α_γ θ_Γ
B
Mean coverage (%)
n=402
n=462
n=573
n=348
α_Γ θ_γ θ_Γ α_γ
C
Mean: 75.0%
Median: 77.8%
Occurrence
Subject coverage (%)
D
Mean: 74.3%
Median: 75.0%
Occurrence
Subject coverage (%)
E
Mean coverage (%)
n=435
n=348
n=474
n=546
α_Γ α_γ θ_γ θ_Γ
F
Mean coverage (%)
n=348
n=435
n=474
n=546
α_γ α_Γ θ_γ θ_Γ
G
Mean: 72.9%
Median: 75.0%
Occurrence
Subject coverage (%)
H
Mean: 72.9%
Median: 75.0%
Occurrence
Subject coverage (%)
I
Mean coverage (%)
n=450
n=387
n=348
n=447
θ_Γ α_γ α_Γ θ_γ
J
Mean coverage (%)
n=387
n=348
n=447
n=450
α_γ α_Γ θ_γ θ_Γ
K
Mean: 74.3%
Median: 75.0%
Occurrence
Subject coverage (%)
L
Mean: 74.2%
Median: 75.0%
Occurrence
Subject coverage (%)
M
Mean coverage (%)
n=162
n=315
n=204
n=321
n=321
γ Γ β θ α
N
Mean coverage (%)
n=315
n=162
n=321
n=204
n=321
Γ γ θ β α
O
Mean: 85.0%
Median: 87.5%
Occurrence
Subject coverage (%)
P
Mean: 85.0%
Median: 86.4%
Occurrence
Subject coverage (%)
Q
Mean coverage (%)
n=246
n=162
n=243
n=249
n=147
Γ β α θ γ
R
Mean coverage (%)
n=246
n=162
n=147
n=243
n=249
Γ γ β α θ
S
Mean: 83.4%
Median: 85.7%
Occurrence
Subject coverage (%)
T
Mean: 84.0%
Median: 85.7%
Occurrence
Subject coverage (%)
U
Mean coverage (%)
n=342
n=249
n=231
n=294
n=270
γ β Γ α θ
V
Mean coverage (%)
n=342
n=249
n=270
n=231
n=294
γ β θ Γ α
W
Mean: 86.1%
Median: 87.5%
Occurrence
Subject coverage (%)
X
Mean: 85.4%
Median: 87.5%
Occurrence
Subject coverage (%)

**Figure S13. Mean and distributions of cross-subject coverage in recognition, encoding, and retrieval tasks for ERPAC and ERPPC analyses**

The figure presents both the mean subject coverage across frequency bands (bar plots, columns 1 and 2) and the overall coverage distributions across all clusters (histograms, columns 3 and 4). The top three rows (panels A–L) display the ERPAC results, while the bottom three rows (panels M–X) show the ERPPC results. Within each method, the rows correspond to the behavioral task: recognition (rows 1 and 4), encoding (rows 2 and 5), and retrieval (rows 3 and 6). For both plot types, the first corresponding column (columns 1 and 3) represents the same condition, and the second corresponding column (columns 2 and 4) represents the other condition. These condition pairs correspond to Novel vs. Non-novel in recognition, Better Encoding vs. Worse Encoding in encoding, and Match vs. Mismatch in retrieval. Subject coverage was defined as the percentage of subjects contributing to a given cluster relative to the total number of subjects available for the corresponding region pair. In the bar plots, bar height indicates the mean subject coverage (%), error bars denote standard deviation, and the text inside each bar indicates the number of clusters (n) in that band. The horizontal red dashed line marks the 80% coverage reference level. In the histograms, vertical dashed lines indicate the mean and median coverage. Across tasks and methods, most clusters were concentrated in the moderate-to-high coverage range, indicating generally robust cross-subject consistency of clustering patterns across behavioral conditions.

### Figure S14

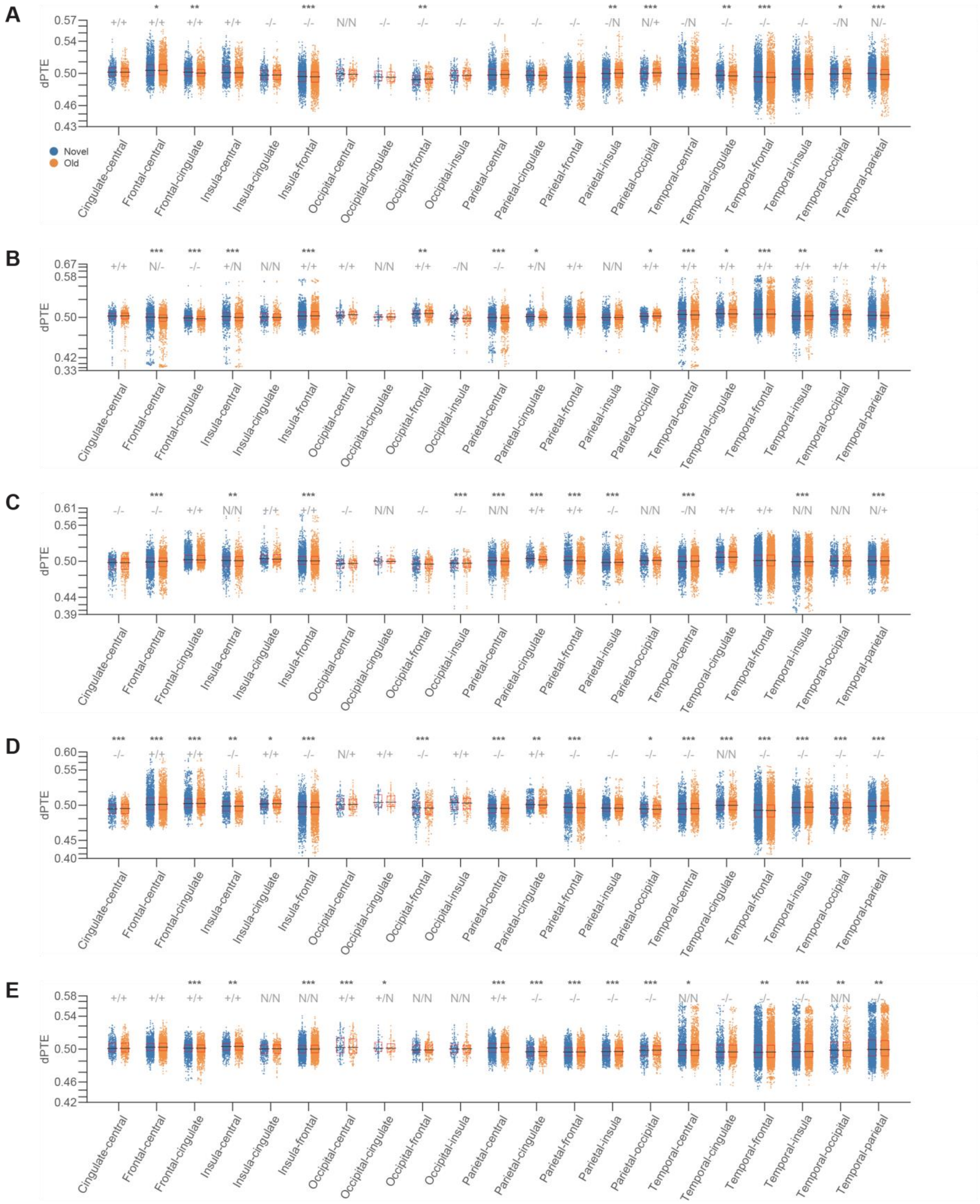
A
B
C
D
E
dPTE
Novel
Old
Cingulate-central
Frontal-central
Frontal-cingulate
Insula-central
Insula-cingulate
Insula-frontal
Occipital-central
Occipital-cingulate
Occipital-frontal
Occipital-insula
Parietal-central
Parietal-cingulate
Parietal-frontal
Parietal-insula
Parietal-occipital
Temporal-central
Temporal-cingulate
Temporal-frontal
Temporal-insula
Temporal-occipital
Temporal-parietal

### Figure S14

**Figure S14. dPTEs difference under memory recognition phase among frequency bands.** dPTE in **(A)** theta, **(B)** alpha, **(C)** beta **(D)** low-gamma and **(E)** high-gamma bands across all subjects (lobe level, novel vs old). Each dot (blue: novel, orange: old) corresponds to the median dPTE of a channel pair in this lobe pair. Asterisk: median dPTEs showed significant difference (Wilcoxon signed rank test, *: $p < 0.05$, **: $p < 0.01$, ***: $p < 0.001$) between novel and old conditions. + means median dPTE was significantly larger than 0.5 (Wilcoxon signed rank test, $p < 0.05$), while – means smaller than 0.5, and N means no significant different compared to 0.5. Black horizontal line: median dPTE of the lobe pair under each condition. Red box: interquartile range ($25^{th}$~$75^{th}$ percentile) of the dPTE distribution for each condition.

### Figure S15

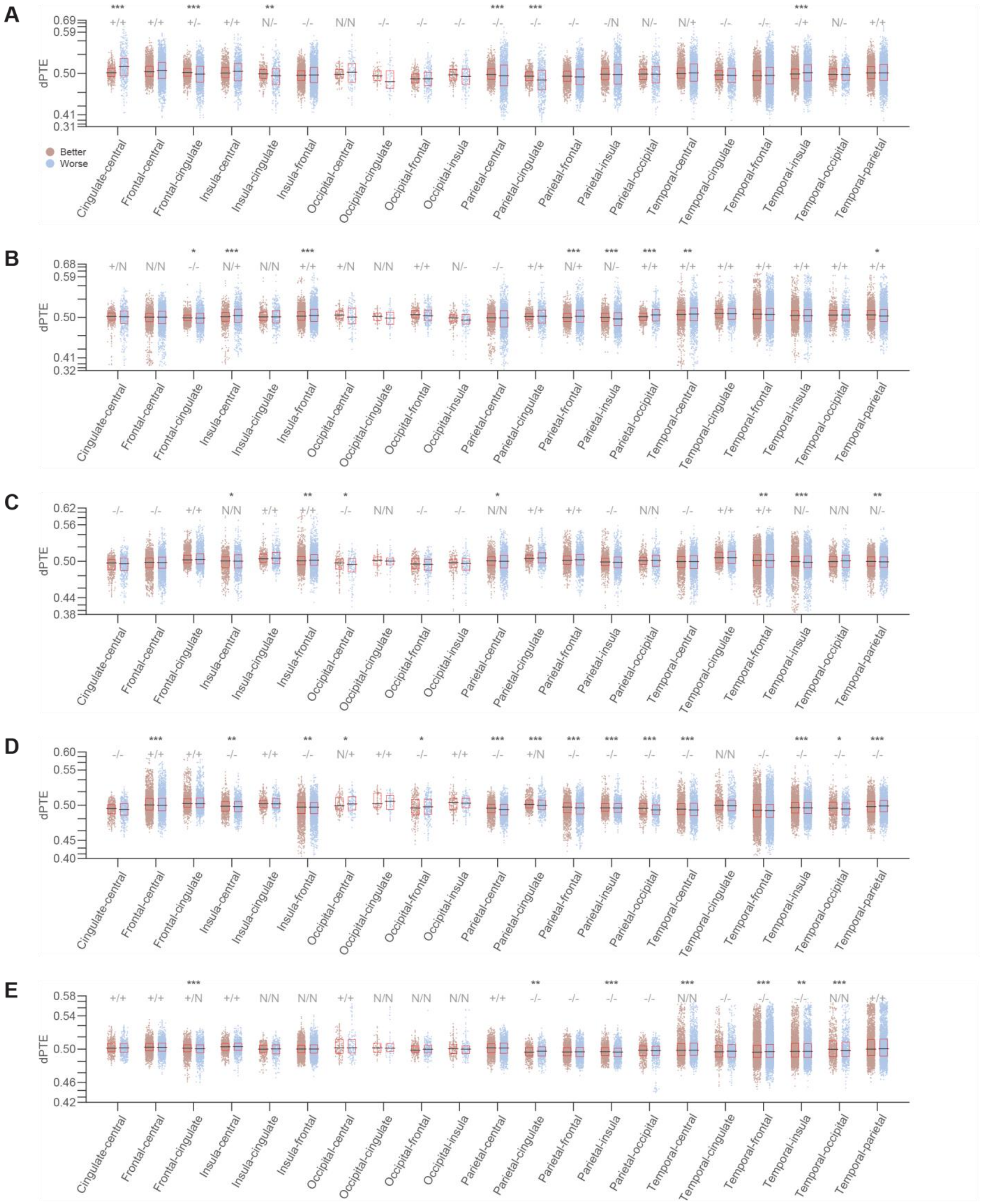

A
B
C
D
E
dPTE
Better
Worse
Cingulate-central
Frontal-central
Frontal-cingulate
Insula-central
Insula-cingulate
Insula-frontal
Occipital-central
Occipital-cingulate
Occipital-frontal
Occipital-insula
Parietal-central
Parietal-cingulate
Parietal-frontal
Parietal-insula
Parietal-occipital
Temporal-central
Temporal-cingulate
Temporal-frontal
Temporal-insula
Temporal-occipital
Temporal-parietal

### Figure S15

**Figure S15. dPTEs difference under memory encoding phase among frequency bands.** dPTE in **(A)** theta, **(B)** alpha, **(C)** beta **(D)** low-gamma and **(E)** high-gamma bands across all subjects (lobe level, better encoding vs worse encoding). Each dot (brown: better encoding, blue: worse encoding) corresponds to the median dPTE of a channel pair in this lobe pair. Asterisk: median dPTEs showed significant difference (Wilcoxon signed rank test, *: $p < 0.05$, **: $p < 0.01$, ***: $p < 0.001$) between better and worse encoding conditions. + means median dPTE was significantly larger than 0.5 (Wilcoxon signed rank test, $p < 0.05$), while – means smaller than 0.5, and N means no significant different compared to 0.5. Black horizontal line: median dPTE of the lobe pair under each condition. Red box: interquartile range ($25^{th}$~$75^{th}$ percentile) of the dPTE distribution for each condition.

### Figure S16

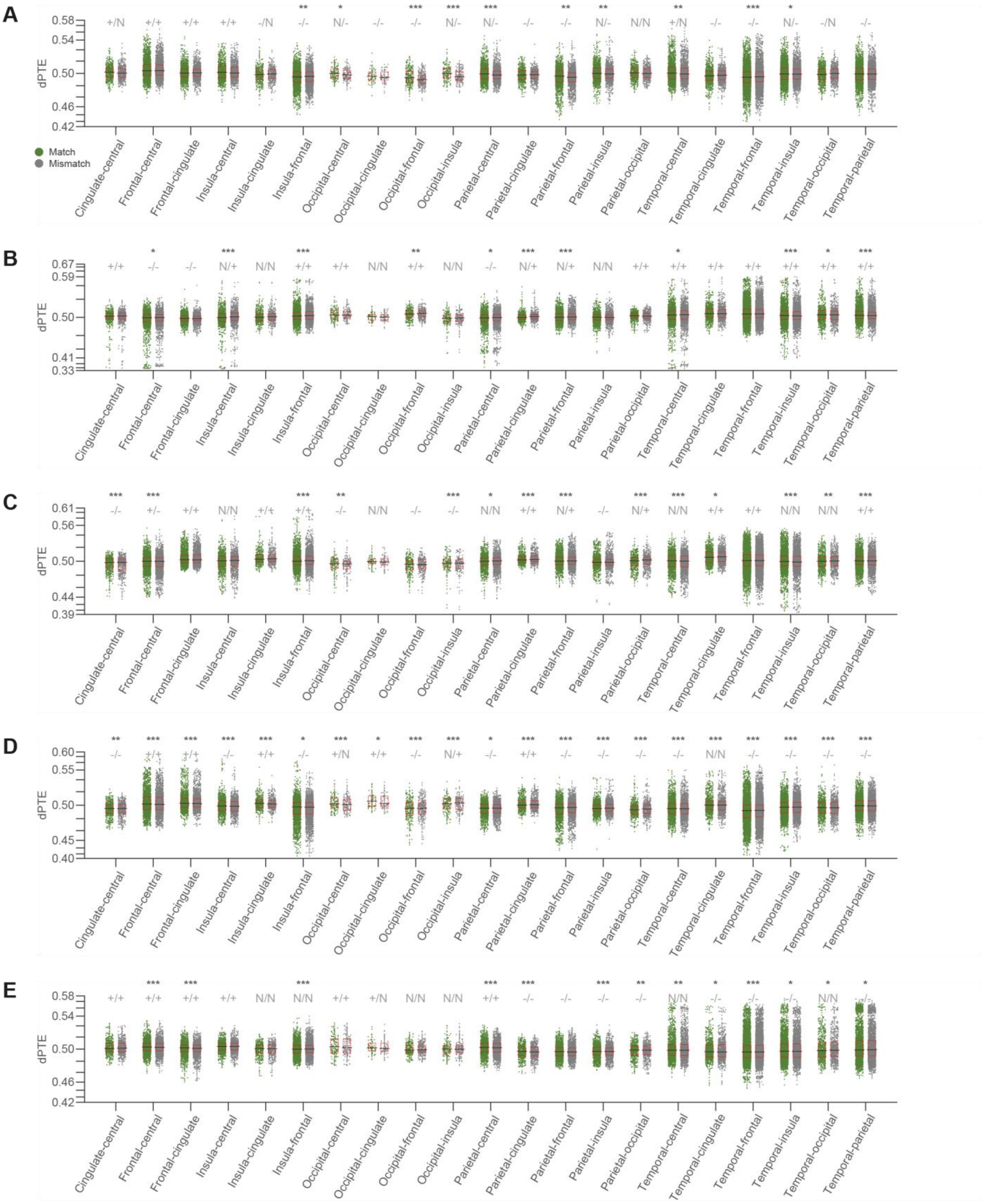
A
B
C
D
E
dPTE
Match
Mismatch
Cingulate-central
Frontal-central
Frontal-cingulate
Insula-central
Insula-cingulate
Insula-frontal
Occipital-central
Occipital-cingulate
Occipital-frontal
Occipital-insula
Parietal-central
Parietal-cingulate
Parietal-frontal
Parietal-insula
Parietal-occipital
Temporal-central
Temporal-cingulate
Temporal-frontal
Temporal-insula
Temporal-occipital
Temporal-parietal

### Figure S16

**Figure S16. dPTEs difference under memory retrieval phase among frequency bands.** dPTE in **(A)** theta, **(B)** alpha, **(C)** beta **(D)** low-gamma and **(E)** high-gamma bands across all subjects (lobe level, match vs mismatch). Each dot (green: match, gray: mismatch) corresponds to the median dPTE of a channel pair in this lobe pair. Asterisk: median dPTEs showed significant difference (Wilcoxon signed rank test, *: $p < 0.05$, **: $p < 0.01$, ***: $p < 0.001$) between match and mismatch conditions. + means median dPTE was significantly larger than 0.5 (Wilcoxon signed rank test, $p < 0.05$), while – means smaller than 0.5, and N means no significant different compared to 0.5. Black horizontal line: median dPTE of the lobe pair under each condition. Red box: interquartile range ($25^{th}$~$75^{th}$ percentile) of the dPTE distribution for each condition.

### Figure S17

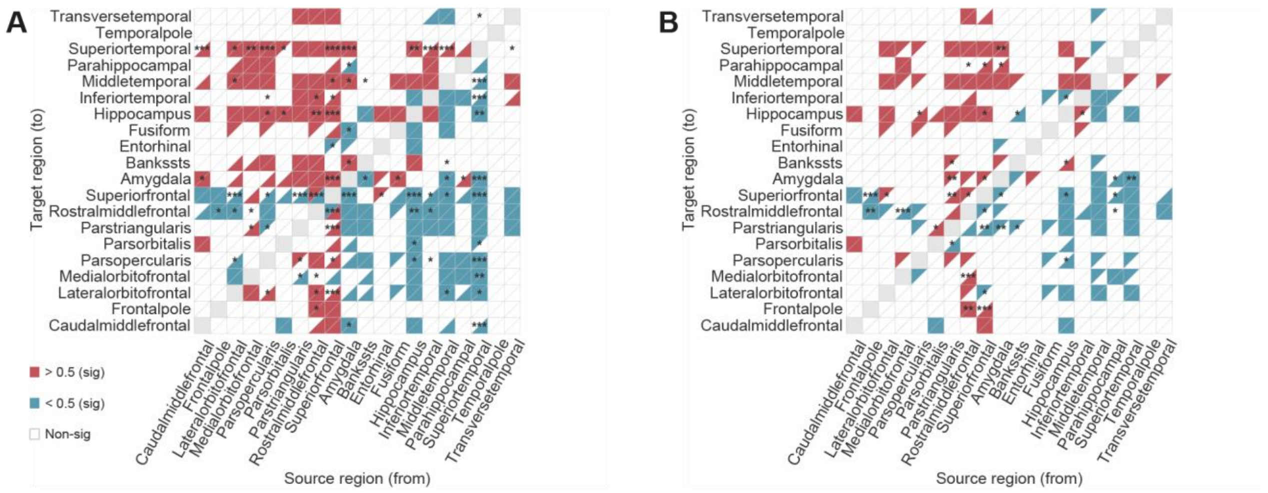


**Figure S17. Region-level dPTEs difference of frontal and temporal lobes in theta band. A.** dPTE difference in theta band for region pairs in frontal and temporal lobes between novel and old conditions. Top-right and bottom-left triangles show whether the dPTEs were significantly larger (red) or lower (blue) than 0.5 under novel and old conditions respectively (Wilcoxon signed rank test, $p < 0.05$). Asterisk: median dPTEs showed significant difference (Wilcoxon signed rank test, *: $p < 0.05$, **: $p < 0.01$, ***: $p < 0.001$) between novel and old conditions. **B.** dPTE difference in theta band for region pairs in frontal and temporal lobes between better encoding and worse encoding conditions. Top-right and bottom-left triangles show whether the dPTEs were significantly larger (red) or lower (blue) than 0.5 under better encoding and worse encoding conditions respectively (Wilcoxon signed rank test, $p < 0.05$). Asterisk: median dPTEs showed significant difference (Wilcoxon signed rank test, *: $p < 0.05$, **: $p < 0.01$, ***: $p < 0.001$) between better encoding and worse encoding conditions.

### Figure S18

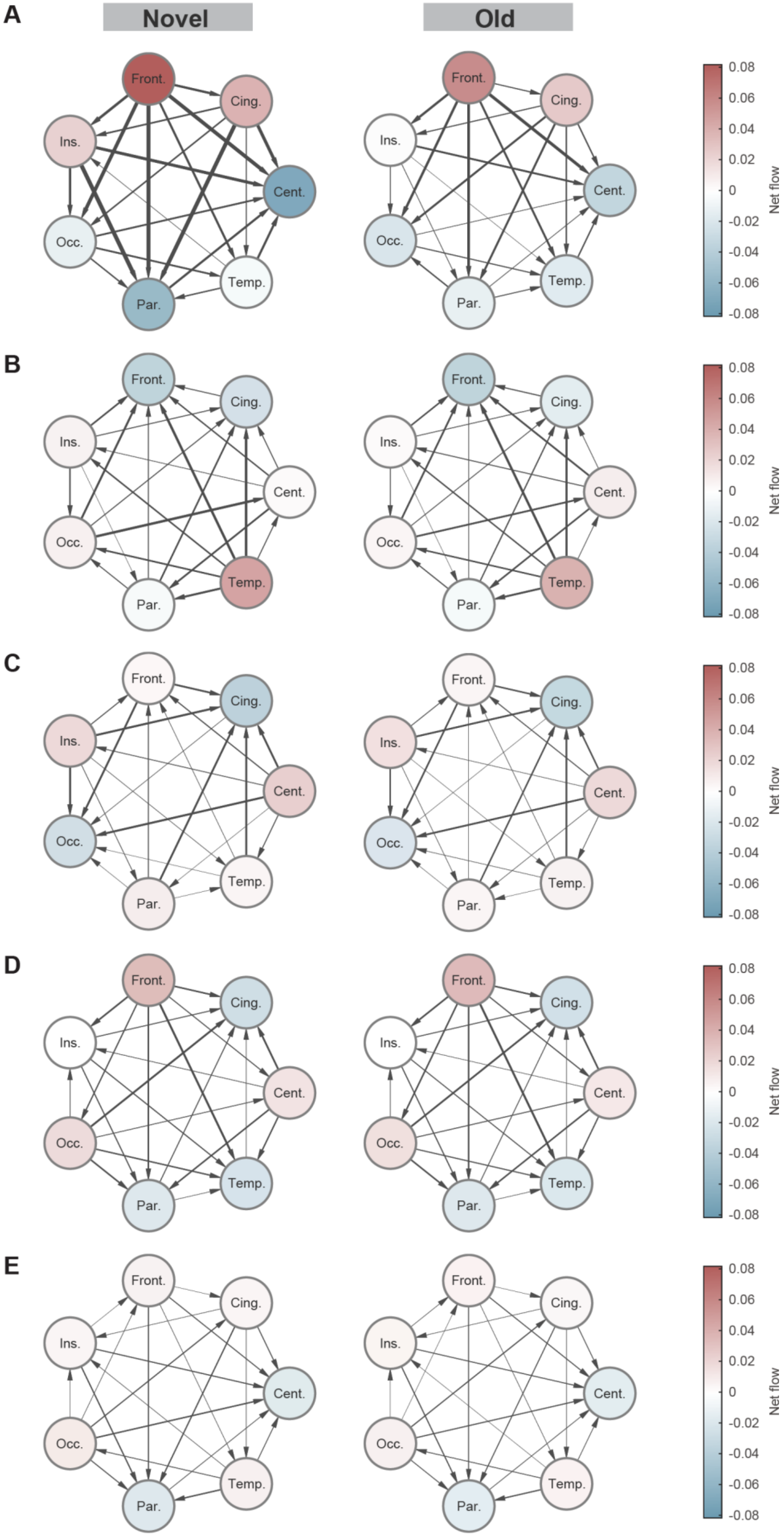


**Figure S18. Networks in memory recognition phase among frequency bands.** Networks in **(A)** theta, **(B)** alpha, **(C)** beta, **(D)** low-gamma and **(E)** high-gamma bands under novel (first column) and old (second column) conditions for all subjects. Net flow: out-strength minus in-strength. Red circle indicates the node acts as an information source (net flow > 0). Blue circle indicates the node acts as an information sink (net flow < 0). Circle size shows the total connectivity of each node (in-degree + out-degree). The thickness of arrows shows the weights between the nodes.

### Figure S19

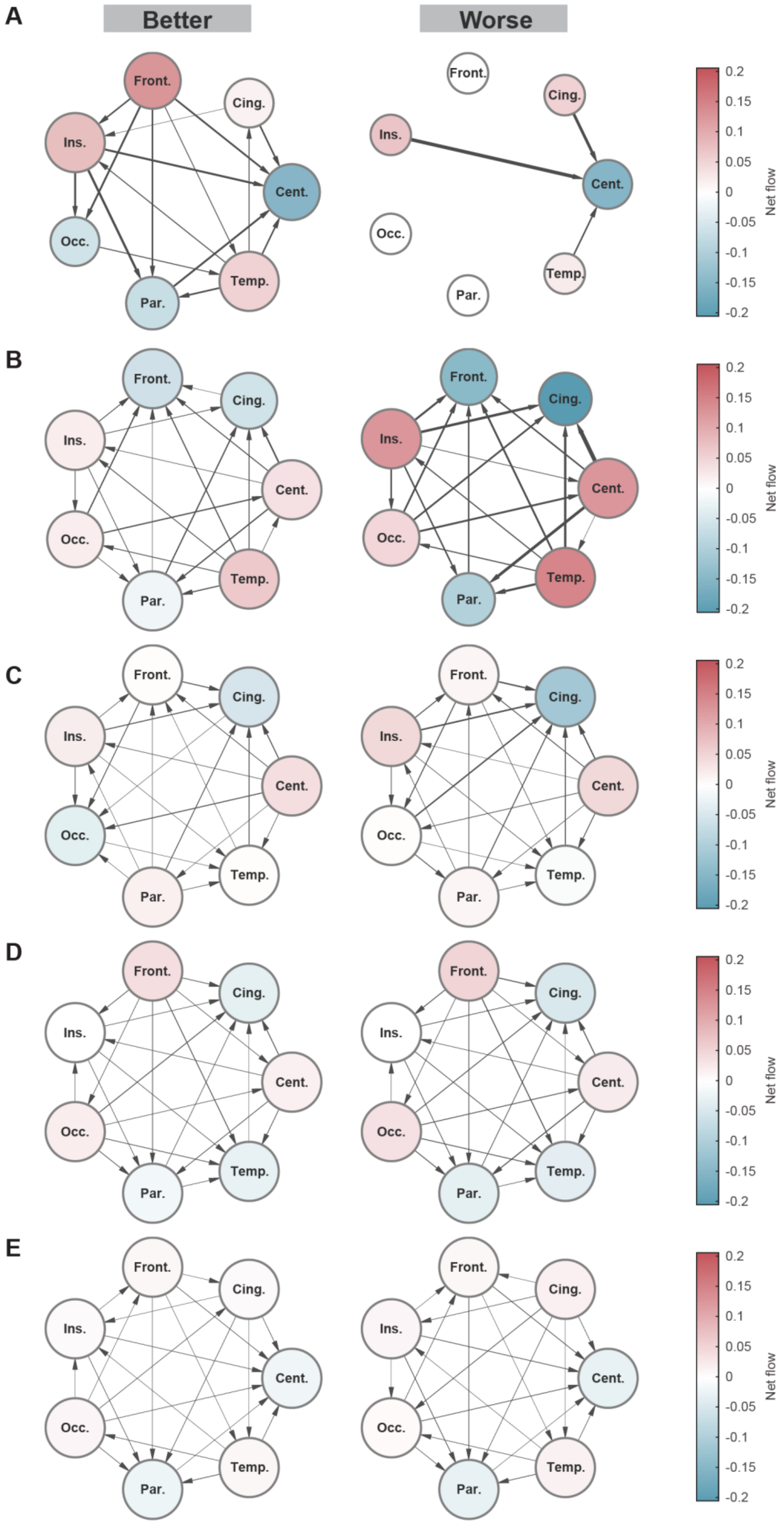


**Figure S19. Networks in memory encoding phase among frequency bands.** Networks in **(A)** theta, **(B)** alpha, **(C)** beta, **(D)** low-gamma and **(E)** high-gamma bands under better encoding (first column) and worse encoding (second column) conditions for all subjects. Net flow: out-strength minus in-strength. Red circle indicates the node acts as an information source (net flow > 0). Blue circle indicates the node acts as an information sink (net flow < 0). Circle size shows the total connectivity of each node (in-degree + out-degree). The thickness of arrows shows the weights between the nodes.

### Figure S20

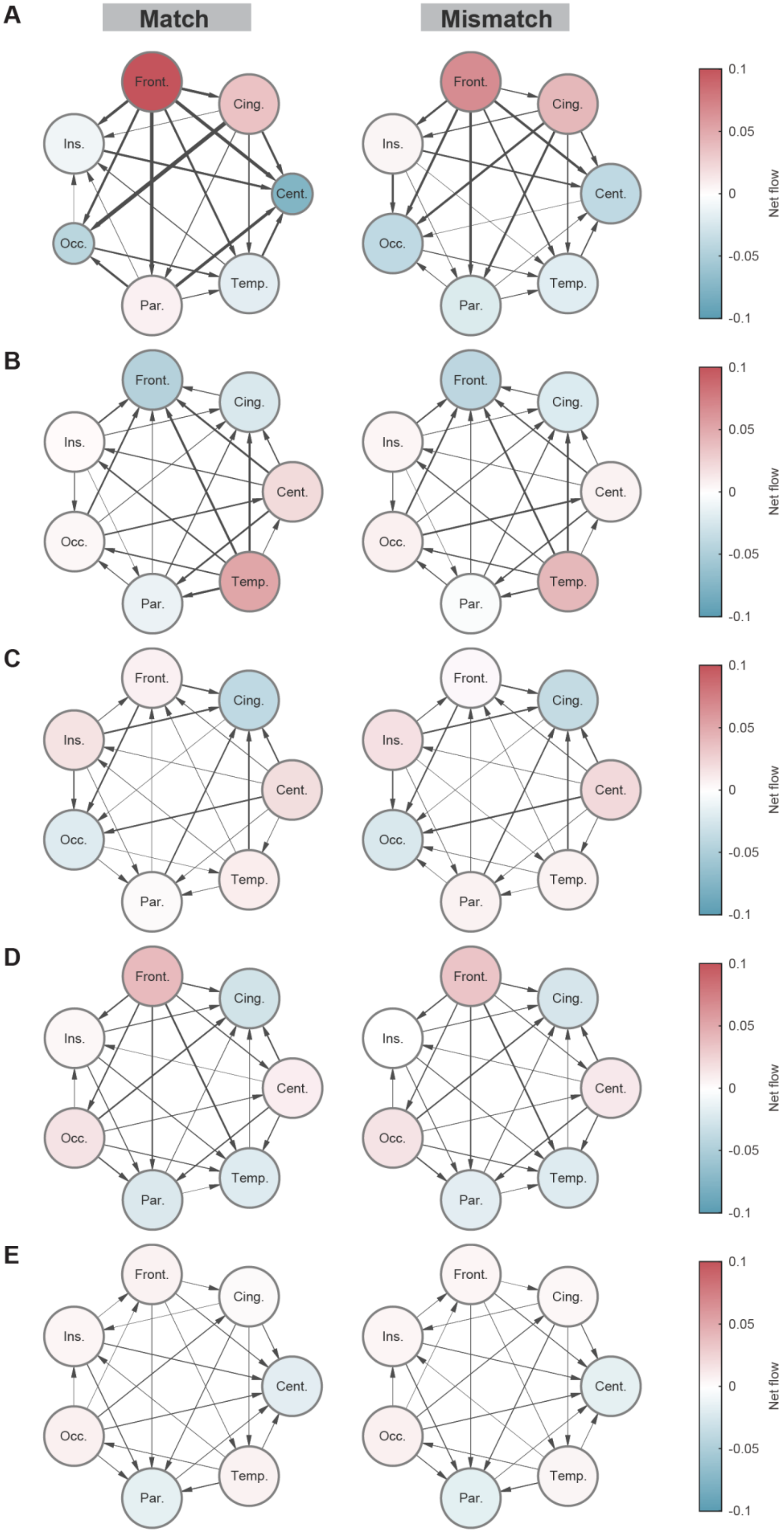


**Figure S20. Networks in memory retrieval phase among frequency bands.** Networks in **(A)** theta, **(B)** alpha, **(C)** beta, **(D)** low-gamma and **(E)** high-gamma bands under match (first column) and mismatch (second column) conditions for all subjects. Net flow: out-strength minus in-strength. Red circle indicates the node acts as an information source (net flow > 0). Blue circle indicates the node acts as an information sink (net flow < 0). Circle size shows the total connectivity of each node (in-degree + out-degree). The thickness of arrows shows the weights between the nodes.

### Figure S21

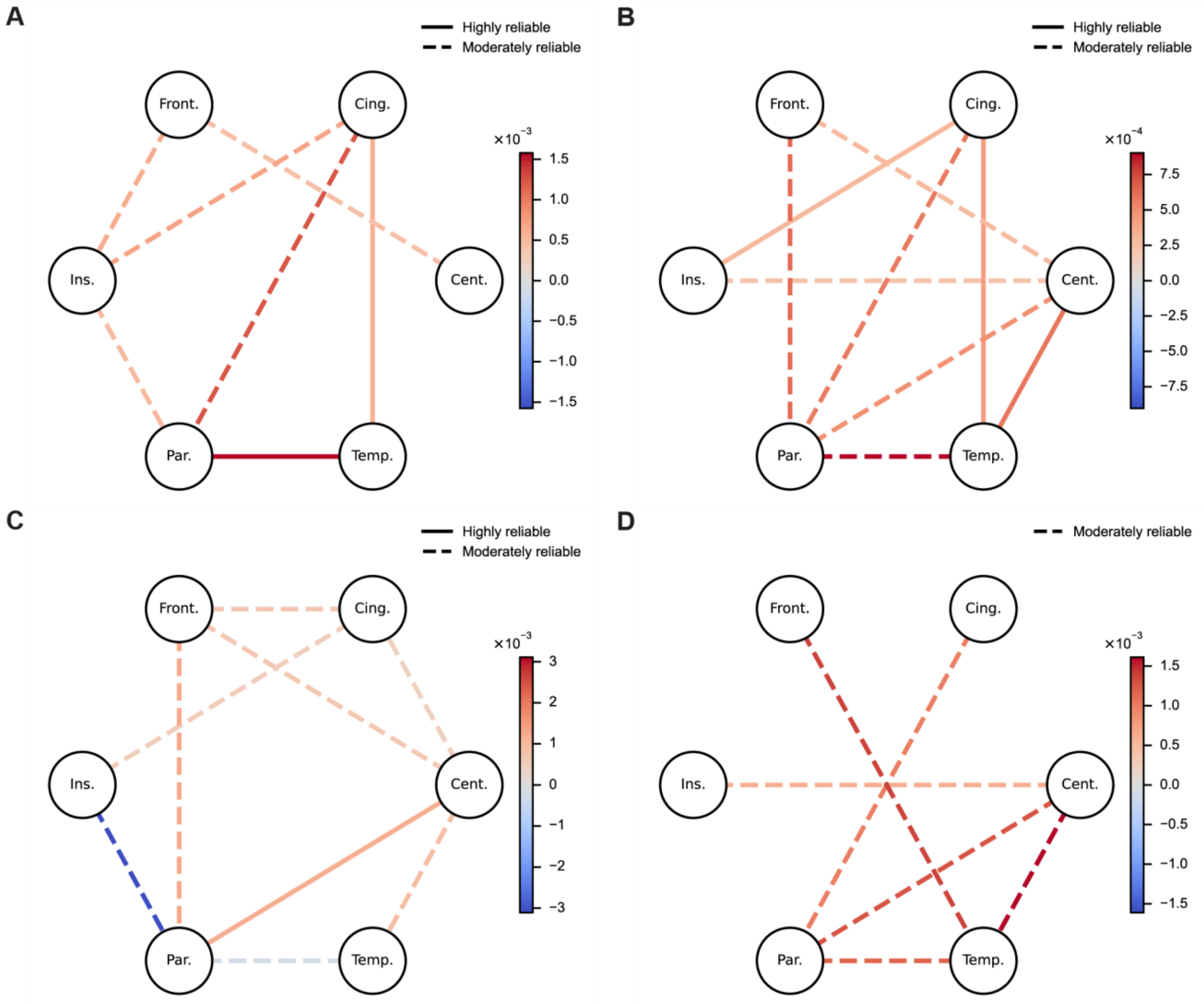


**Figure S21. Lobe-level coherence contrast within frequency bands.** Lobe-level coherence contrast in the (**A**) theta, (**B**) beta, (**C**) low-gamma, and (**D**) high-gamma bands between the two memory-related network configurations (late-dominant minus early-dominant). Node labels denote cortical lobes; edge color encodes mean Δcoherence, and line style reflects cross-subject reliability (solid: highly reliable; dashed: moderately reliable).
